# AlphaFold-Multimer Modelling of Linked nAChR Subunits Challenges Concatemer Design Assumptions

**DOI:** 10.1101/2025.10.02.679753

**Authors:** Hanna M Sahlström, Lucien Rufener, Kristian Høy Horsberg, Anouk Sarr, Tor Einar Horsberg, Marit J Bakke

## Abstract

Nicotinic acetylcholine receptors (nAChRs) are well described in vertebrates, yet less studied in arthropods, and the subunit stoichiometry of heteromeric arthropod nAChRs remains unresolved. This study combined a computational and experimental approach to predict and validate the stoichiometries of two heteromeric nAChRs from the parasitic arthropod *Lepeophtheirus salmonis*. AlphaFold2- and AlphaFold-Multimer-based modelling, supported by multiple sequence alignment and functional expression in *Xenopus laevis* oocytes, identified the most likely stoichiometries for two compositions of subunits previously described, named Lsa-nAChR1 and Lsa-nAChR2. For receptor Lsa-nAChR1, the highest scoring stoichiometry was α1b1α2α2b2. Lsa-nAChR2 exhibited three possible stoichiometries, confirmed by both computational modelling and experiments. These are α3β2α3β1β2, α3β1α3β1β2, and α3α3α3β1β2. All stoichiometries are written from a counterclockwise, extracellular orientation. Strikingly, structural modelling also suggested that linker flexibility in concatemer constructs may allow a novel conformation with a different subunit between the linked subunits, referred to here as “wedging”.

These results indicate that the use of flexible linker sequences does not reliably enforce subunit position or assembly directionality, as shown here for nAChRs. These findings challenge the assumption that linked concatemers unambiguously dictate receptor stoichiometry. Thus, interpretations of concatemer-based studies, on nAChRs and other receptor systems, may warrant careful reevaluation.

## Introduction

Nicotinic acetylcholine receptors (nAChRs) are ligand-activated pentameric cation channels found in the central and peripheral nervous system, muscles, and other tissues of many organisms. They mediate fast synaptic transmission in both vertebrates and invertebrates and play a vital role in relaying signals across the nervous system. In mammals, nAChRs have been extensively studied and serve as therapeutic targets for conditions such as nicotine addiction, Alzheimer’s disease, schizophrenia, Parkinson’s disease, and chronic pain^1,2^. Each receptor consists of five separate subunits that form an ion channel. These subunits are categorized as α, β, γ, δ, and ε. The ion channel is permeable to sodium, potassium, and in some cases, calcium ions. Channel opening occurs when the natural ligand acetylcholine (ACh) binds at the interface between an α-subunit and either another α-subunit or a non-α-subunit^3^. Functional heteromeric nAChRs with resolved structures contain at least two α-subunits and, consequently, two ACh binding sites^4^.

In contrast to mammals, much less is known about nAChRs in insects and other arthropods. In these organisms, nAChRs are the primary targets for important control agents, such as neonicotinoids. One such neonicotinoid insecticide is imidacloprid, which was recently approved for use against the salmon louse (*Lepeophtheirus salmonis*), a major parasite in salmonid aquaculture ^5–7^. Despite their pharmacological importance, the subunit composition and stoichiometry of arthropod nAChRs remain poorly characterized. This limits the current understanding of their functions and hinders the development of selective modulators.

Although the subunit composition and function of arthropod native nAChRs are largely unknown, several nucleotide sequences of invertebrate nAChR subunits have been identified. Genome sequencing of an increasing number of arthropods has provided insights into these receptors. One experimental method to study their function is to inject heterologous capped RNA (cRNA) into unfertilized oocytes from *Xenopus laevis* to express membrane proteins on the oocyte surface. The functions of the expressed channels can subsequently be studied using electrophysiological techniques^8^. Despite the significant role of these receptors in the control of pests and parasites, and ongoing attempts to clone them for decades^9^, no functional nAChRs assembled from non-chimeric arthropod α- and β-subunits had been successfully expressed in any *ex vivo* system until 2020.

In 2020, the successful expression of two heterologous nAChRs from *L. salmonis* into *Xenopus laevis* oocytes was published^10^. These two receptors, named Lsa-nAChR1 and Lsa-nAChR2, shared two β-subunits but differed in which α-subunits they consisted of^10^. Functional formation of Lsa-nAChR1 consisted of α1, α2, β1 and β2 subunits, while α3, β1, and β2 subunits were necessary to form functional Lsa-nAChR2 receptors. Since these receptors are pentamers, some of the subunits must appear multiple times in both Lsa-nAChR1 and Lsa-nAChR2. There is a strong indication that one of the two unidentified subunits in Lsa-nAChR2 must be an α3, since nAChRs are expected to have at least two ACh-binding sites^4^. These binding sites are found at the interface between an α-subunit and an adjacent subunit^4^. However, the identity of the subunits could not be determined by the experimental set-up published in 2020.

Different approaches have been employed to determine the exact subunit order and composition in heteromeric receptors. For instance, both site-directed mutagenesis of a conserved residue combined with electrophysiology^11^, and fluorescently tagged antibodies against individual subunits have been used to determine the stoichiometry of the GABA_A_ receptor^12^. Fluorescently labelled subunits have also been used to determine the stoichiometry of several nAChR-receptors^13^. Furthermore, direct subunit counting was used to determine the ratio of subunits in a glycine receptor^14^.

Another experimental approach widely adopted in subsequent studies of receptor composition and stoichiometries is the technique where different subunits are linked together with a synthetic peptide, forcing specific subunit arrangements)^15,16^. These “linked” subunits are called concatemers and allow for functional testing of pre-determined subunit order in systems such as *Xenopus laevis* oocytes. This is done by injecting different subunit sequences in one long cRNA fragment, separated by the sequence encoding the peptide linker. Concatemers have been a powerful tool in determining receptor structure and have contributed to the characterization of numerous ion channels in both vertebrate and invertebrate systems^15,17–22^. Widely used, concatemer-based approaches rely on the assumption that linker sequences rigidly constrain subunit positioning. This assumption has rarely been tested directly and is widely based on the expectation that the “lowest-energy” configuration will position the linked subunits next to each other. The structural consequences of linker flexibility on subunit arrangement remain poorly understood. A further limitation of this approach is that a “brute force” attempt to construct all theoretically possible stoichiometries of a receptor is costly and labour-intensive. For example, for the salmon louse Lsa-nAChR1 which consists of α1, α2, β1 and β2 subunits^10^, there are 48 theoretical stoichiometries that need to be functionally verified *in vitro*. Thus, an *in silico* approach to predict the most likely stoichiometry using bioinformatic tools could greatly reduce laboratory workload.

Recent advances in protein structure prediction, particularly the development of AlphaFold2 and AlphaFold-Multimer, have enabled the accurate modelling of protein complexes based solely on amino acid sequence information^23,24^. These tools offer a computational alternative to experimentally intensive and expensive methods such as protein crystallography. AlphaFold has been widely recognized as a paradigm-shifting breakthrough in structural biology, enabling high-confidence predictions of protein structures at an unprecedented scale^25^.

The overall aim of the current study was to explore a bioinformatic pipeline to predict the most likely stoichiometry of Lsa-nAChR1 and Lsa-nAChR2. This involved first ranking model scores *in silico*, using tools such as AlphaFold2, AlphaFold-Multimer, ZRANK, and Prodigy, and then comparing the highest-scoring models to *ex vivo* functional tests of linked subunit compositions in the *Xenopus laevis* oocyte model.

## Materials and methods

### Annotation of Subunit Positions

Subunit compositions, for example α3β2α3β1β2, are always written without hyphens separating individual subunits in the counterclockwise order of assembly and viewed from the extracellular side. The principal component is defined as the subunit located to the left at a dimer interface in the external top-down view of the pentamer. It contributes to the principal face of the binding site, including the C-loop that clasps over acetylcholine upon binding.^3^ The subunit located to the right at a dimer interface is defined as the complementary component. Linked (concatenated) subunits are indicated with hyphens, for example α3-β2.

### Structural Modelling of Lsa-nAChRs using AlphaFold2 and Scored-Based Evaluation

To predict the three-dimensional structures of theoretically possible heteropentamers of the *L. salmonis* Lsa-nAChR1 and Lsa-nAChR2 receptors, standalone versions of AlphaFold2 and AlphaFold-Multimer^23,24^ were run under Linux Ubuntu 22.04.5 on a Lenovo ThinkStation P720 equipped with an NVIDIA RTX A4000 graphics card, using default settings. The amino acid sequences of the subunits were downloaded from GenBank (α1: QHU23855, α2: QHU23856, α3: QHU23857, β1: QHU23859, β2: QHU23860). Signal peptides, predicted by the online software DeepTMHMM^26^, were removed from the downloaded fasta files before modelling. The final input fasta files can be found at https://gitlab.com/hansahls/nachr-stoichiometry-repo/-/tree/main/Output_Alphafold2_Computer/Alphafold2_Produced_Pentamers.

AlphaFold2 and AlphaFold-Multimer predict a number of possible stoichiometries for the multimers and assess and rank the quality of the different predictions for each given combination of subunits (for example subunits α1,α1,α2,β1, and β2 for the Lsa*-*nAChR1). The scores and rankings are given for these predictions; however, this assessment is not directly comparable to other subunit combinations (for example subunits α1, α2, α2, β1, and β2). The score is a combination of how well the whole complex fits a “true” structure, and the accuracy of the relative positions of the subunits in the prediction. Therefore, two additional tools were used to assess the docking quality between the predicted dimers of the subunits. After modelling each of the possible pentamers of Lsa-nAChR1 and Lsa-nAChR2, each pdb-pentamer file was split into five adjacent dimer pdb-files. Each dimer was then tested for protein-protein interactions using standalone versions of the bioinformatic packages ZRANK^27^, and PRODIGY^28^. For each package, the individual scores for the five dimers in a pentamer were summed to obtain an overall score for the pentamer. To allow for cross-comparison across the different score types by AlphaFold2, ZRANK, and PRODIGY, robust scaling was applied to each score type. Specifically, each score was normalized by subtracting the median and dividing by the interquartile range (IQR) of all values within that score type. The resulting scaled values were then summed across all three score types to generate a composite “robust score” for each pentamer model. This normalization approach, based on the RobustScaler method^29,30^ is referred to here as “total robust scaling”, and allowed for comparative ranking of the produced pentamer models while preserving relative variation within each metric. The generated pdb models, scores, and scripts can be found at https://gitlab.com/hansahls/nachr-stoichiometry-repo.

Robust scaling was chosen over standard z-score normalization due to ZRANK and PRODIGY scores having skewed distributions and the presence of outliers. This violates the assumption of normality required for mean-based scaling. Using the median and IQR provides a more robust and interpretable scale for combining metrics derived from heterogeneous sources, as well as this method being robust to outliers. ^29–33^

The illustrations of three-dimensional protein structures were made with the software ChimeraX^34^

### Multiple Sequence Alignment of Ligand Binding Loops Across Species

To assess the conservation of functionally important ligand-binding residues, amino acid sequences from *L. salmonis* nAChR subunits were aligned with homologous sequences from *Apis mellifera*, *Aedes albopictus*, *Drosophila melanogaster*, and *Homo sapiens*. Sequences were retrieved from NCBI and aligned using ClustalW implemented in UGENE^35^. Ligand-binding loop regions (A-F) were defined according to the structural classifications described by Grutter and Changeux ^36^. Residues previously reported to directly participate in ligand binding were annotated manually.

### Experimental determination of stoichiometry

*In silico* docking of individual subunits predicts the exact stoichiometry, but this needs to be verified experimentally. In the experimental section, concatemers were injected and expressed in *X. laevis* oocytes. For each nAChR subunit, a “START” and a “STOP” construct were generated using specially designed primers (Supplementary Material C). For “START” constructs, the stop codon was deleted, a flexible amino acid linker of variable length (Supplementary Material D) was inserted, as well as XmaI, EagI, AgeI and XhoI restriction sites. For “STOP” constructs, the signal peptide was deleted and replaced with a stretch of restriction sites (NheI, AgeI, AvrII, AflII, and ApaI). The amino acid linker connected the C-terminus of the first subunit to the N-terminus of the second subunit. In addition, “CASSETTE” constructs were also amplified: those constructs lack the signal peptides as well as the stop codon. NheI and AgeI restriction sites were added at the 5’-end while ApaI and XhoI were added at the 3’-end. Each construct was cloned into the pT7-TS expression vector and sequenced to ensure that the Open Reading Frame (ORF) was intact.

Using this approach, all possible dimers of Lsa-nAChR1 and Lsa-nAChR2 can be theoretically generated. All the generated concatemers are listed in Table 3 and Table 4. The cRNA generated for these concatemer was injected into *X. laevis* oocytes together with the cRNA encoding free subunit(s), other concatemers, and necessary chaperone proteins (RIC-3, UNC-50, and UNC-74) to determine which of them formed functional nAChRs ^10^. All the injected concatemer and subunit combinations can be found in Table 3 and Table 4.

### Electrophysiology

Capped cRNAs were synthesized (T7 mMessage mMachine kit, Ambion, Austin, TX, USA) from linearized vectors containing different concatemers or unlinked/free subunits according to the manufacturer’s protocol. cRNA samples were stored at -80°C until use. Oocytes were ordered from Ecocyte (Germany) and upon arrivalwere transferred individually to a 96-well plate into sterile filtered Barth solution containing: NaCl (88 mM), KCl (1 mM), NaHCO_3_ (2.4 mM), HEPES (10 mM, pH 7.5), MgSO_4_ ˑ 7 H_2_O (0.82 mM), Ca(NO_3_)_2_ ˑ 4 H_2_O (0.33 mM), CaCl_2_ ˑ 6H_2_O (0.41 mM), at pH 7.4, and supplemented with 20 μg/ml of kanamycin, 100 U/ml benzylpenicillin and 100 μg/ml streptomycin. Following this, the oocytes were microinjected using a Roboinject automatic injection system (Multi Channel Systems, Reutlingen, Germany) with 15-25 nl of cRNA solution per oocyte (30-300 ng/μl per subunit). Subsequently, the oocytes were incubated at 18°C. Recordings were made 3-5 days after cRNA injection.

The recordings were carried out using a two-electrode voltage-clamp setup (HiClamp, MultiChannel Systems). In short, two electrodes in glass capillaries filled with 3M KCl were impaled into the oocyte. The oocyte’s membrane potential was then maintained at -80 mV throughout the experiment and the currents evoked by 100µM ACh were recorded as the compensating current needed to keep the clamped potential. Data was captured at 100Hz and filtered at 20Hz.

### Post-experimental Evaluation

All wet-lab tested concatemer and free subunit combinations were compared to the full set of theoretically possible stoichiometries which included at least two α-subunits. This included 48 possible configurations for Lsa-nAChR1 and 16 for Lsa-nAChR2.

### Post-Experimental Modelling of Linked Subunits in AlphaFold2

Following the experimental analysis, AlphaFold-Multimer was used to model linked pentameric subunit constructs. These constructs were identical to the experimentally tested concatemers and were included to explore the structural outcomes arising from linking subunits. Modelling parameters were applied as described above, with the only modification being the inclusion of variable peptide linkers connecting the C-terminus of the first subunit to the N-terminus of the second subunit. It was assumed that the first subunit would be arranged as the principal subunit in the concatemer and the second subunit as the complementary subunit. The linkers used are listed in Supplementary Material D, and the fasta files for the concatemers can be found at https://gitlab.com/hansahls/nachr-stoichiometry-repo/-/tree/12eb499d4108698f7dfa9844ef09a9d11cfafd9e/Output_Alphafold2_Computer/Linker_Models/fasta/dimers_trimers_linked.

### Data Availability

All data generated or analysed in this study are available in public repositories. The AlphaFold2-predicted receptor models, docking results, and scoring outputs are accessible via GitLab at:https://gitlab.com/hansahls/nachr-stoichiometry-repo

This includes:

- Predicted pentamer structures for Lsa-nAChR1 and Lsa-nAChR2
- Dimer interface files and scores
- Sequence alignments and conservation analyses
- Supplementary electrophysiological data

### Code Availability

All scripts used for AlphaFold2 model scoring, robust scaling, and visualization are available in the same GitLab repository: https://gitlab.com/hansahls/nachr-stoichiometry-repo

## Results

### Structural Modelling and Evaluation of Lsa-nAChR1 Pentamers and Dimers

For Lsa-nAChR1, AlphaFold2 predicted 30 unique stoichiometries out of the 48 theoretical stoichiometry configurations. All 16 possible dimer combinations were represented in these models.

The ranks of the five best scores according to AlphaFold2, PRODIGY, and ZRANK for each dimer summed to pentamer are presented in Table 1. All model rankings and scores are included in Supplementary Material E-F, and can also be found in https://gitlab.com/hansahls/nachr-stoichiometry-repo/R_scripts

**Table 1.** Order of the five best pentamers containing the subunits α1, α2, β1 and β2 in Lsa-nAChR1 from *L. salmonis*. The pentamers were scored with AlphaFold2. Dimer interfaces were scored using ZRANK and PRODIGY, and summed to give a total score per model. The Rank Sum represents the sum of the individual ranks for AlphaFold2, PRODIGY and ZRANK.

| Stoichiometry | AlphaFold Score (iptm+ptm) | PRODIGY Score | ZRANK Score | Rank Sum |
| --- | --- | --- | --- | --- |
| $\alpha 1\beta 1\alpha 2\alpha 2\beta 2$ | 0.765639013 | -121.3 | 1883.29 | 18 |
| $\alpha 1\alpha 2\beta 2\beta 1\beta 2$ | 0.775139217 | -113.1 | 1901.12 | 18 |
| $\alpha 1\beta 2\alpha 2\beta 1\beta 2$ | 0.77587734 | -109.4 | 1909.74 | 18.5 |
| $\alpha 1\beta 1\beta 2\beta 2\alpha 2$ | 0.752849716 | -117.3 | 1809.23 | 42 |
| $\alpha 1\beta 1\beta 2\alpha 2\beta 1$ | 0.747960286 | -118 | 1957.08 | 47 |

### Structural Modelling and Evaluation of Lsa-nAChR2 Pentamers and Dimers

For Lsa-nAChR2, AlphaFold2 predicted 12 unique stoichiometries out of the 16 theoretical stoichiometry configurations. Within these models, all possible 9 dimer combinations were represented. The ranks of the six best scores according to AlphaFold, PRODIGY, and ZRANK for each dimer summed to pentamer are presented in Table 2. All model rankings are included in Supplementary Material G-H.

**Table 2.**
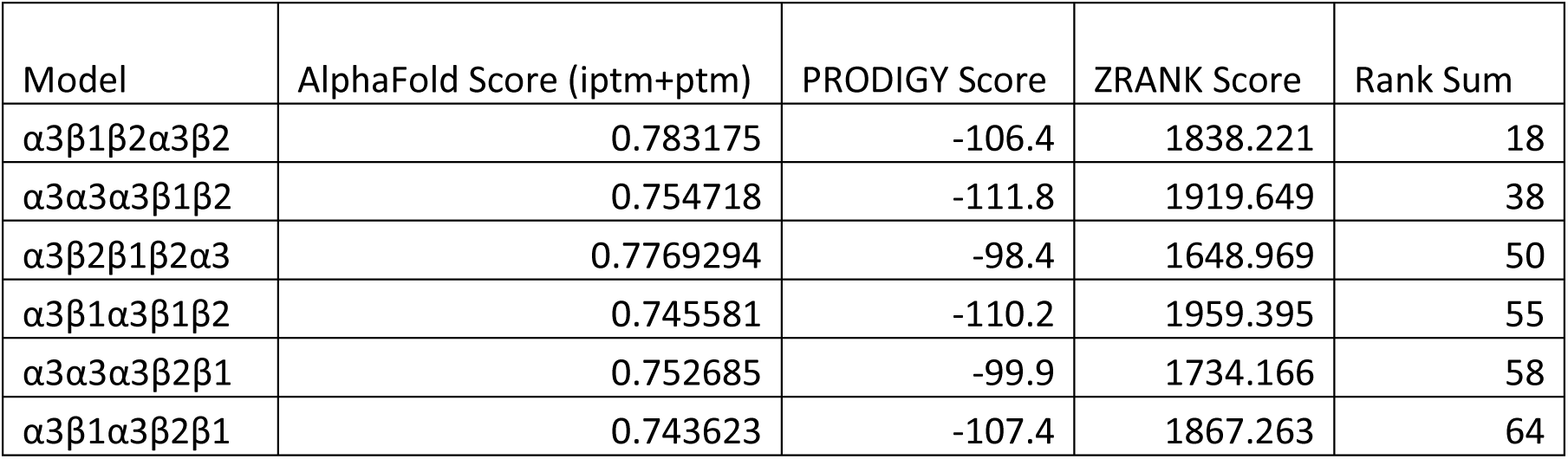
Ranks of the six best pentamers containing the subunits α3, β1 and β2 in Lsa-nAChR2 from *L. salmonis*. The pentamers were scored with AlphaFold2. Dimer interfaces were scored using ZRANK and PRODIGY, then summed to give a total score per model. The rank represents the sum of the individual ranks for AlphaFold2, PRODIGY, and ZRANK. Ranks are skipped until the next model variation is observed.

| Model | AlphaFold Score (iptm+ptm) | PRODIGY Score | ZRANK Score | Rank Sum |
| --- | --- | --- | --- | --- |
| $\alpha 3\beta 1\beta 2\alpha 3\beta 2$ | 0.783175 | -106.4 | 1838.221 | 18 |
| $\alpha 3\alpha 3\alpha 3\beta 1\beta 2$ | 0.754718 | -111.8 | 1919.649 | 38 |
| $\alpha 3\beta 2\beta 1\beta 2\alpha 3$ | 0.7769294 | -98.4 | 1648.969 | 50 |
| $\alpha 3\beta 1\alpha 3\beta 1\beta 2$ | 0.745581 | -110.2 | 1959.395 | 55 |
| $\alpha 3\alpha 3\alpha 3\beta 2\beta 1$ | 0.752685 | -99.9 | 1734.166 | 58 |
| $\alpha 3\beta 1\alpha 3\beta 2\beta 1$ | 0.743623 | -107.4 | 1867.263 | 64 |

### Combined Visualization of Total Robust Scores across Stoichiometries

To summarize the score distributions of all modelled stoichiometries, the total robust scores for the Lsa-nAChR1 and Lsa-nAChR2 pentamer models were plotted as a boxplot using ggplot2 in R, as shwon in Figure 1. Each dot represents a single pentamer model, grouped according to its subunit order and composition. The boxplots highlight variation in “best fit” within each stoichiometry, while the number of dots indicates how frequently each stoichiometry was generated. For Lsa-nAChR2, the stoichiometry α3β1β2α3β2 exhibited both the highest median score and the greatest number of modelled pentamers.

**Figure 1:**
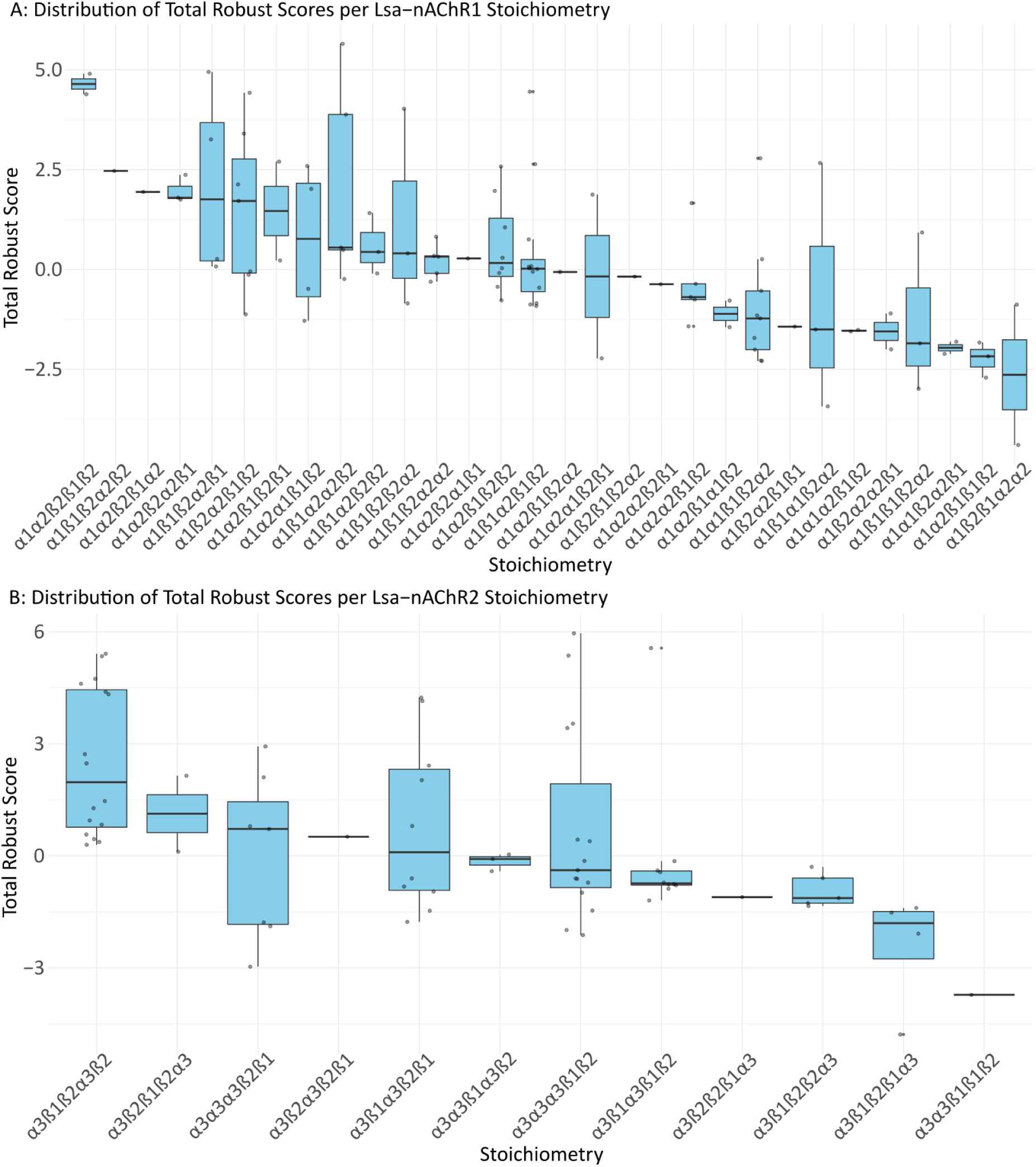
Total robust score distribution across modelled stoichiometries for Lsa-nAChR1 and Lsa-nAChR2. Each boxplot shows the distribution of the total robust scores for pentamer models grouped by stoichiometry. For Lsa-nAChR1, 100 pentamer models were analysed; for Lsa-nAChR2, 75 pentamer models were analysed. Individual dots represent the total robust scores of individual pentamer models, allowing visualization of both the distribution and the number of models generated per stoichiometry. The stoichiometries are arranged from left to right in order of decreasing median total robust score. Scores were calculated by summing the robust-scaled values from AlphaFold2, ZRANK, and PRODIGY, as described in the Methods section.

Notably, two other stoichiometry groups exhibited models with high robust scores: a β1-rich configuration (α3 β1α3β2β1), and a triple-α3 configuration (α3α3α3β1β2). These three stoichiometries were consistently ranked among the highest in both raw and scaled scoring metrics. For Lsa-nAChR1, a wider range of unique stoichiometries were generated, which reduced the number of models per configuration. Among these, the stoichiometry α1β1α2α2β2 had the highest individual scoring pentamer model, whereas α1α2β2β1β2 showed the highest median score.

### Cross-Species Comparison of Ligand Binding Loop Conservation in Lsa-nAChRs

To evaluate conservation of ligand-binding residues in salmon lice nAChRs, multiple sequence alignments of α- and β-subunits were analysed from *L. salmonis*, *D. melanogaster*, *A. mellifera*, and *H. sapiens*, focusing on loop regions A-F as defined by Grutter et al, 2001^36^.

Loops A, B, and C which comprise the principal component subunit, presented with high sequence conservation among invertebrate α-subunits. This was particularly noticeable at positions corresponding to key residues in the human receptor: W149 and Y151 in Loop A, and Y190, C192, C193, and Y198 in Loop C. Notably, Lsa-α2 displayed a phenylalanine (F) substitution at the position aligned with Y93 in Loop B, whereas all other α-subunits retained a conserved Loop B region. The β-subunits displayed greater variability in these principal component residues, consistent with their complementary roles in ligand binding. These alignment regions are shown in Figure 2.

**Figure 2:**
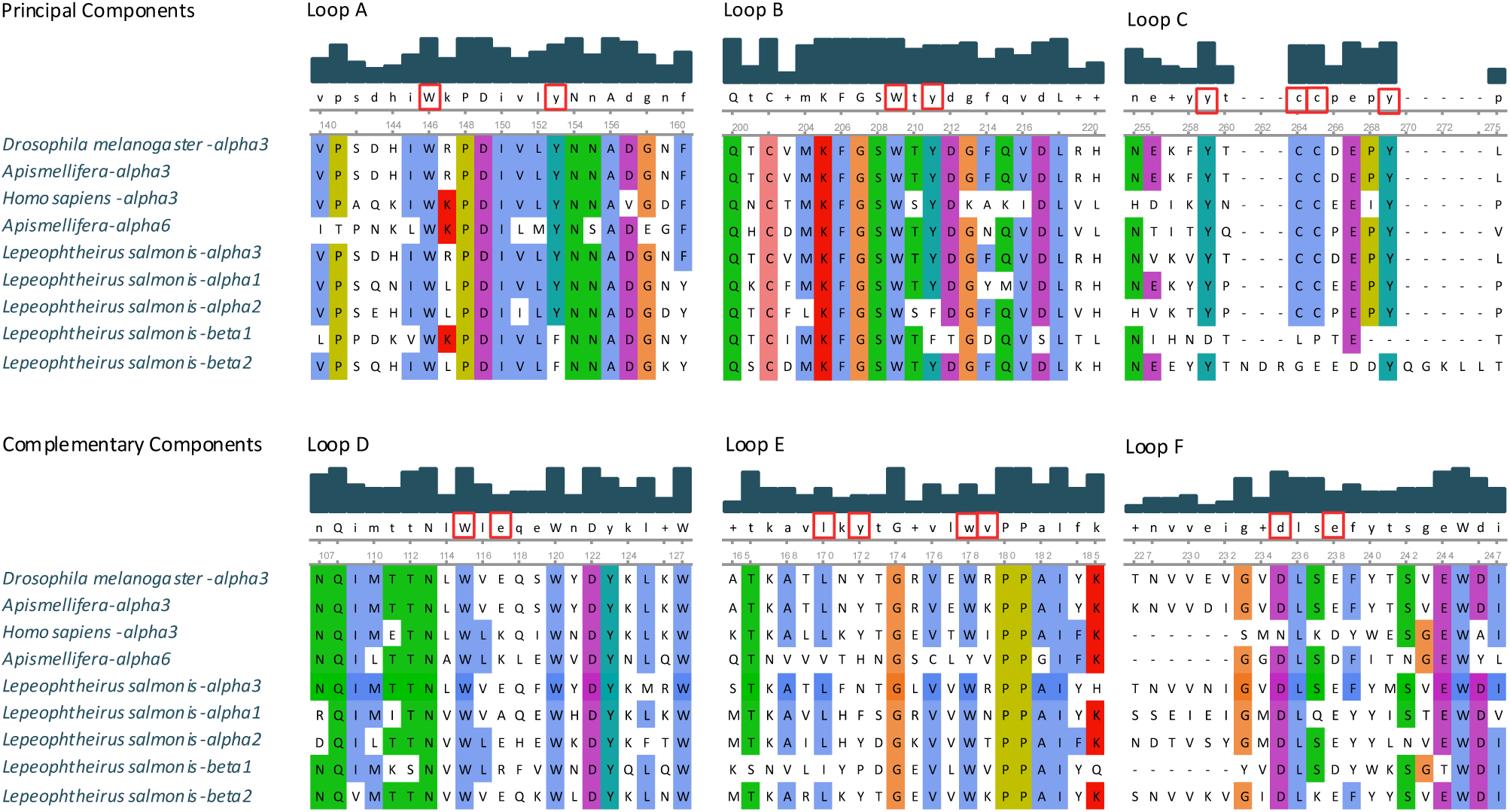
Sequence alignment of nAChR ligand-binding loops in vertebrates and invertebrates. Amino acid sequences from α- and β-subunits of vertebrate and invertebrate species were aligned to determine the conservation of key residues that form binding loops A-F. Histogram bars represent per column conservation, and residues boxed in red represent functionally important conserved positions, based on alignment with human α3 subunits. The species and subunit identities are listed to the left. Colouring reflects amino acid identity using the Clustal colour scheme.

Loops D-F displayed greater sequence variability across both α- and β-subunits in invertebrates. Subsequent positions, particularly in Loop E, were only loosely conserved and were absent in certain species entirely. Conversely, residues W55 and E57 forming Loop D were fully conserved among α3-subunits from *D. melanogaster*, *A. mellifera*, and *L. salmonis*. In *L. salmonis*, these residues are also conserved in the α2- and β2-subunits. Partial conservation of Loop F was seen in *D. melanogaster* α3, *A. mellifera* α3, and the *L. salmonis* α1, α2, α3, and β2 subunits. Interestingly, the β1-subunit of *L. salmonis* lacked conserved residues in all three complementary loops.

### Experimental determination of stoichiometry

For Lsa-nAChR1, testing concatemers and free subunits in *Xenopus laevis* oocytes showed that only three of thirteen constructs produced a functional receptor. The combinations tested are listed in Table 3. For Lsa-nAChR2, the test of concatemers and free subunits revealed that eight constructs injected together with free subunits gave rise to a functional receptor, while eight did not. The combinations tested are listed in Table 4.

**Table 3.** Concatemers tested experimentally in the *Xenopus* oocyte model for Lsa-nAChR1. The concatemers were either tested together with free subunits, concatemers, or both. Concatemers linked with a polypeptide chain are indicated as two subunits with a hyphen (e.g. α2-α2). The possible orientations were determined from AlphaFold2 modelling of the concatemers as dimers or trimers.

| Receptor | Concatemers | Free Subunits | Functional |
| --- | --- | --- | --- |
| R1 | α2-α2 | α1; β1; β2 | YES |
| R1 | β2-α1 | α2; β1 | YES |
| R1 | $\alpha 2\text{-}\alpha 2\text{-}\beta 2$ | $\alpha 1; \beta 1$ | YES |
| R1 | $\alpha 2\text{-}\alpha 1$ | $\beta 1; \beta 2$ | NO |
| R1 | $\alpha 2\text{-}\beta 1$ | $\alpha 1; \beta 2$ | NO |
| R1 | $\alpha 2\text{-}\beta 2$ | $\alpha 1; \beta 1$ | NO |
| R1 | $\beta 1\text{-}\alpha 1$ | $\alpha 2; \beta 2$ | NO |
| R1 | $\beta 1\text{-}\alpha 2$ | $\alpha 1; \beta 2$ | NO |
| R1 | $\beta 1\text{-}\beta 1$ | $\alpha 1; \alpha 2; \beta 2$ | NO |
| R1 | $\beta 1\text{-}\beta 2$ | $\alpha 1; \alpha 2$ | NO |
| R1 | $\beta 2\text{-}\alpha 2$ | $\alpha 1; \beta 1$ | NO |
| R1 | $\beta 2\text{-}\beta 1$ | $\alpha 1; \alpha 2$ | NO |
| R1 | $\beta 2\text{-}\beta 2$ | $\alpha 1; \alpha 2; \beta 1$ | NO |

**Table 4.**
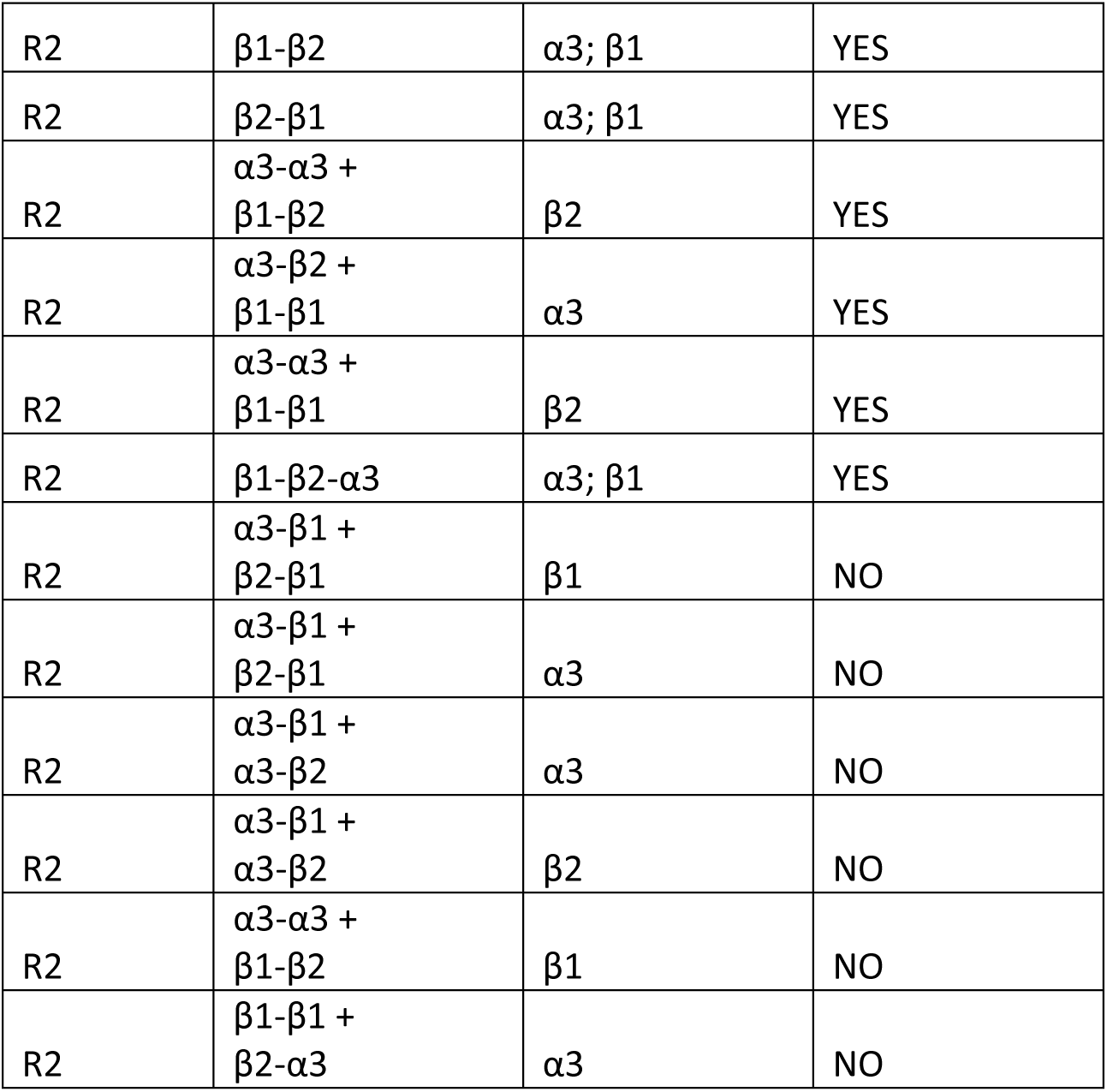
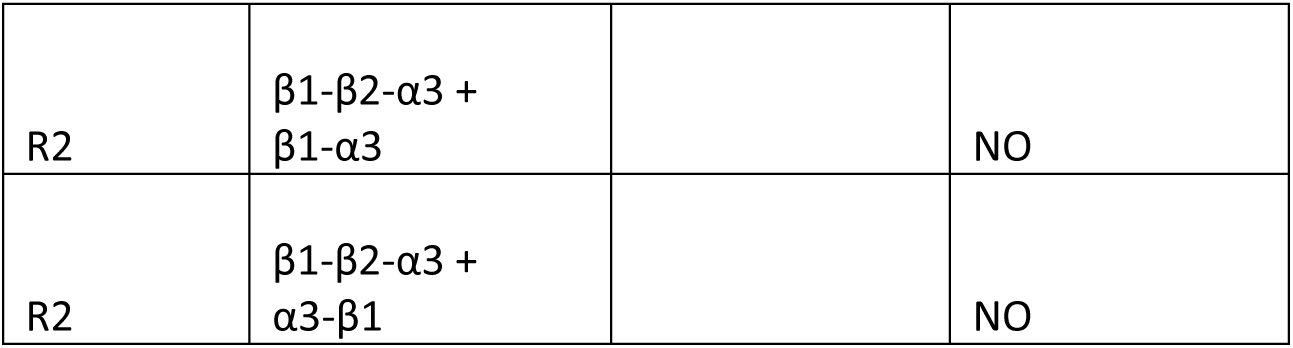
Concatemers tested experimentally In the *Xenopus* oocyte model for Lsa-nAChR2. The concatemers were either tested together with free subunits, or concatemers, or both. Concatemers linked with a polypeptide chain are indicated as two subunits with a hyphen (e.g. β1-β2). The possible orientations were determined from AlphaFold2 modelling of the concatemers as dimers or trimers.

|  |  |  |  |
| --- | --- | --- | --- |
| R2 | $\beta 1\text{-}\beta 2$ | $\alpha 3; \beta 1$ | YES |
| R2 | $\beta 2\text{-}\beta 1$ | $\alpha 3; \beta 1$ | YES |
| R2 | $\alpha 3\text{-}\alpha 3 + \beta 1\text{-}\beta 2$ | $\beta 2$ | YES |
| R2 | $\alpha 3\text{-}\beta 2 + \beta 1\text{-}\beta 1$ | $\alpha 3$ | YES |
| R2 | $\alpha 3\text{-}\alpha 3 + \beta 1\text{-}\beta 1$ | $\beta 2$ | YES |
| R2 | $\beta 1\text{-}\beta 2\text{-}\alpha 3$ | $\alpha 3; \beta 1$ | YES |
| R2 | $\alpha 3\text{-}\beta 1 + \beta 2\text{-}\beta 1$ | $\beta 1$ | NO |
| R2 | $\alpha 3\text{-}\beta 1 + \beta 2\text{-}\beta 1$ | $\alpha 3$ | NO |
| R2 | $\alpha 3\text{-}\beta 1 + \alpha 3\text{-}\beta 2$ | $\alpha 3$ | NO |
| R2 | $\alpha 3\text{-}\beta 1 + \alpha 3\text{-}\beta 2$ | $\beta 2$ | NO |
| R2 | $\alpha 3\text{-}\alpha 3 + \beta 1\text{-}\beta 2$ | $\beta 1$ | NO |
| R2 | $\beta 1\text{-}\beta 1 + \beta 2\text{-}\alpha 3$ | $\alpha 3$ | NO |
| R2 | $\beta 1\text{-}\beta 2\text{-}\alpha 3 + \beta 1\text{-}\alpha 3$ | | NO |
| R2 | $\beta 1\text{-}\beta 2\text{-}\alpha 3 + \alpha 3\text{-}\beta 1$ | | NO |

For Lsa-nAChR1, functional receptor formation was only observed when an α2-α2 concatemer was used, strongly constraining the possible functional stoichiometries. This supports a composition of α1, α2, α2, β1, and β2, aligning with the highest total robust score and ranked AlphaFold2 model.

*Ex vivo* testing of concatemer constructs for Lsa-nAChR2 functionally validated the presence of three separate stoichiometries capable of forming functional receptors. The successful expression of the concatemer combination α3-α3, β1-β2 with a free β2 subunit supports the existence of a β2-rich configuration. Likewise, the success of α3-β2 with β1-β1 with a free α3 subunit, as well as the success of the combination α3-α3, β1-β1, with a free β2 subunit supports a β1-rich composition. Finally, the functional expression of a β1-β1 concatemer with a free α3 subunit supports an α3-rich configuration.

These data indicate that Lsa-nAChR2 is functionally competent in at least three distinct subunit compositions, aligning closely with the high-scoring configurations observed *in silico*.

### Structural Outcomes of Linked Subunit Modelling in AlphaFold2 and AlphaFold-Multimer

Although the subunit composition of each receptor could be identified through *ex vivo* experiments, the exact order of these subunits remained unclear. Further modelling was performed using AlphaFold2. These models were based on the exact subunit combinations and linker sequences used in the wet-lab concatemer constructs. Specifically, dimeric and trimeric concatemers were modelled with peptide linkers and with free subunit combinations identical to those used in the experimental electrophysiology analysis, and predicted as full pentamers. This approach aimed to determine whether the subunits forming the concatemers only could be arranged in one fixed configuration, or if alternative structural arrangements due to linker flexibility were possible. Using this method, several novel concatemer variations were identified. The concatemers were classified into four different groups. A: *Counterclockwise*, B: *Clockwise*, C: *Wedge Type 1*, and D: *Wedge Type 2*, presented as simplified illustrations in Figure 3.

**Figure 3:**
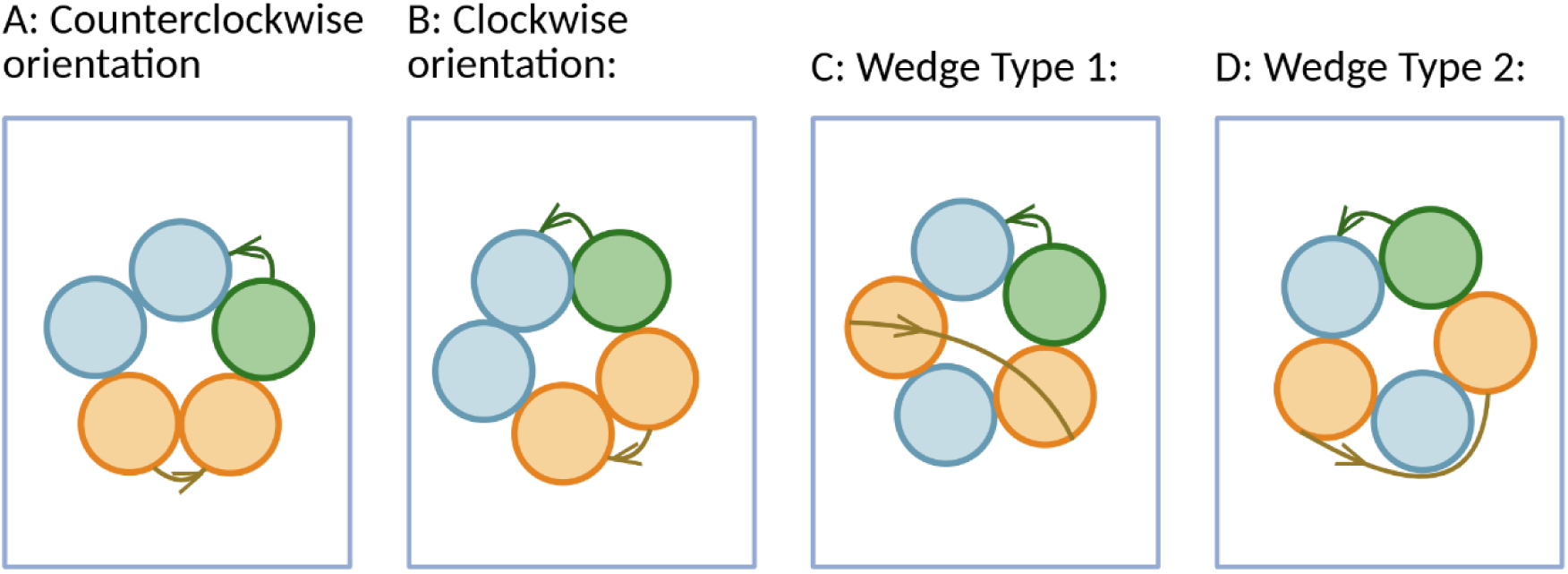
Panels A-D depict simplified illustrations of Figure 4, showing four observed structural variations of the linker between the orange subunits: (A) *Counterclockwise*, (B) *Clockwise*, (C) *Wedge Type 1*, and (D) *Wedge Type 2*. The *Counterclockwise* arrangement (A) shows the linked subunits arranged adjacently in a counterclockwise order. The *Clockwise* orientation (B) shows the linked concatemers arranged in a clockwise order. *Wedge Type 1* shows the linker crossing over the pore of the channel with another subunit fitting in between the linked subunits, while *Wedge Type 2* shows the linker on the outside of the receptor, also with another subunit fitting in between the linked subunits.

Figure 4 presents representative AlphaFold2-predicted concatemers. These were modelled using the same linker sequences and subunit combinations tested experimentally in the oocyte electrophysiology analysis. The results presented here are for the combination α3-α3, β1-β2 and a free β2 subunit; however the full list is available in Supplementary Material I. These structures included both expected and unexpected conformations. Examples of unexpected assemblies are defined and referenced here as *Wedge Type 1* and *Wedge Type 2*. *Wedge Types 1* and *2* were observed when a free subunit wedged in between a linked dimer pair. *Wedge Type 1* is classified as when the linker crosses over the pore, possibly blocking it. *Wedge Type 2* is classified as when the linker wraps around the outside of the protein complex.

**Figure 4:**
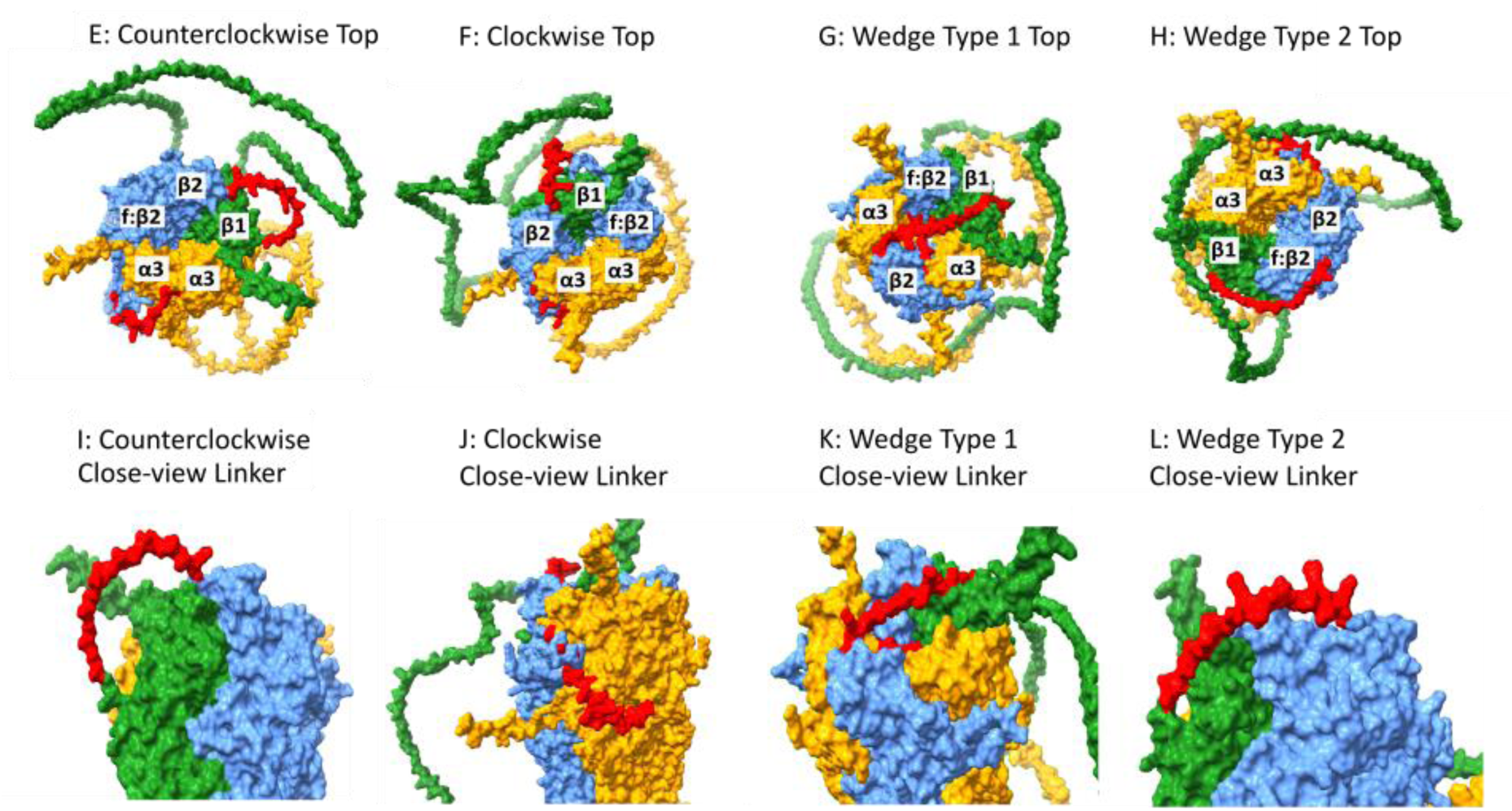
Panels E-L depict the results of AlphaFold2-predicted conformational variability of linked pentamers from the experiment using two linked dimers, α3-α3 and β1-β2, and one free β2 subunit. Panels E-H show top-down (extracellular) views of the predicted assemblies for each conformation. Panels I-L provide close-ups of the linkers in each type of arrangement. Subunits are coloured by type (α3 = yellow, β1= green, β2 = blue), and linker sequences are highlighted in red. The unlinked free subunit is marked as *f:β2*.

Table 5 provides an overview of a couple representative concatemer input configurations found in Supplementary Material I, and their corresponding AlphaFold2-predicted outputs. Each linker is categorised as *Counterclockwise* (CC), *Clockwise* (Cl), *Wedge Type 1* (WdT1) or *Wedge Type 2* (WdT2). Model confidence scores are included, with values above 0.75 are used as a cut-off to indicate high structural reliability. While there is no benchmark set for iptm+ptm scores, previous studies have used similar metrics^37,38^. Notably, the Lsa-nAChR2 model with the highest prediction score corresponds to the subunit arrangement α3β1β2α3β2, the same configuration predicted by AlphaFold2 when the subunits were modeled as free, non-concatenated components. All concatemer models producing this configuration require wedged structural rearrangements to achieve this subunit order. In contrast, this was not the case for Lsa-nAChR1, where the input concatemer permitted an adjacent concatemer arrangement to achieve the highest scoring configuration: α1β1α2α2β2.

**Table 5:**
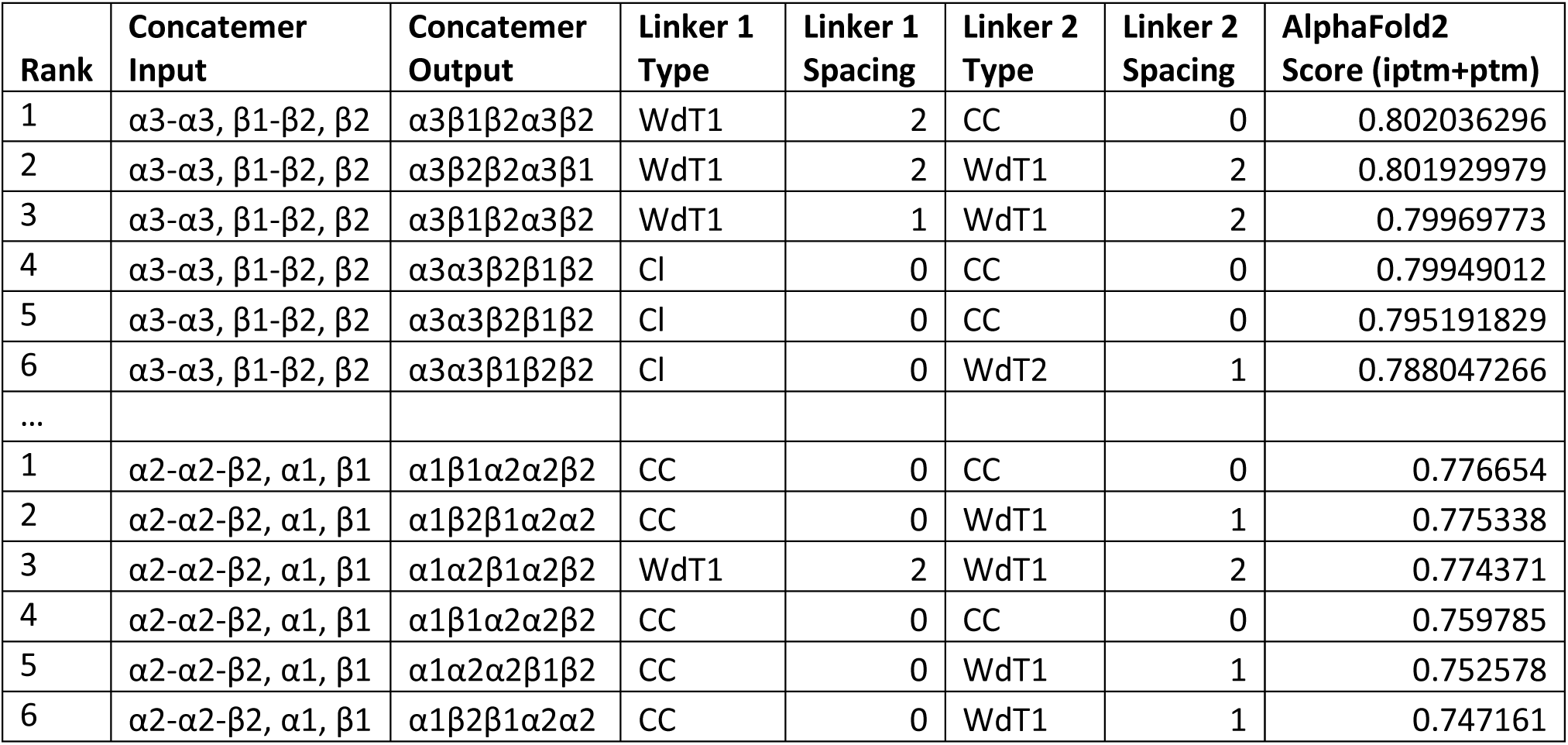
Ranks of AlphaFold2-predicted linked concatemers modelled based on experimental concatemers that produced positive signals in *Xenopus* oocyte tests. Commas indicate the end of one modelled concatemer input, and the beginning of the next input (concatemer or free subunit). The full table can be found in Supplementary Material I. In the input column, linked subunits are indicated with hyphens (e.g. β1-β2), while free subunits are written without hyphens (e.g. …, β2). Each linker position is specified as CC (counterclockwise), Cl (clockwise), WdT1 (Wedge Type 1), or WdT2 (Wedge Type 2). The number of subunits between the linked subunits in the counterclockwise direction is specified in the Linker Spacing columns.

| Rank | Concatemer Input | Concatemer Output | Linker 1 Type | Linker 1 Spacing | Linker 2 Type | Linker 2 Spacing | AlphaFold2 Score (iptm+ptm) |
| --- | --- | --- | --- | --- | --- | --- | --- |
| 1 | $\alpha 3\text{-}\alpha 3, \beta 1\text{-}\beta 2, \beta 2$ | $\alpha 3\beta 1\beta 2\alpha 3\beta 2$ | WdT1 | 2 | CC | 0 | 0.802036296 |
| 2 | $\alpha 3\text{-}\alpha 3, \beta 1\text{-}\beta 2, \beta 2$ | $\alpha 3\beta 2\beta 2\alpha 3\beta 1$ | WdT1 | 2 | WdT1 | 2 | 0.801929979 |
| 3 | $\alpha 3\text{-}\alpha 3, \beta 1\text{-}\beta 2, \beta 2$ | $\alpha 3\beta 1\beta 2\alpha 3\beta 2$ | WdT1 | 1 | WdT1 | 2 | 0.79969773 |
| 4 | $\alpha 3\text{-}\alpha 3, \beta 1\text{-}\beta 2, \beta 2$ | $\alpha 3\alpha 3\beta 2\beta 1\beta 2$ | CI | 0 | CC | 0 | 0.79949012 |
| 5 | $\alpha 3\text{-}\alpha 3, \beta 1\text{-}\beta 2, \beta 2$ | $\alpha 3\alpha 3\beta 2\beta 1\beta 2$ | CI | 0 | CC | 0 | 0.795191829 |
| 6 | $\alpha 3\text{-}\alpha 3, \beta 1\text{-}\beta 2, \beta 2$ | $\alpha 3\alpha 3\beta 1\beta 2\beta 2$ | CI | 0 | WdT2 | 1 | 0.788047266 |
| ... |  |  |  |  |  |  |  |
| 1 | $\alpha 2\text{-}\alpha 2\text{-}\beta 2, \alpha 1, \beta 1$ | $\alpha 1\beta 1\alpha 2\alpha 2\beta 2$ | CC | 0 | CC | 0 | 0.776654 |
| 2 | $\alpha 2\text{-}\alpha 2\text{-}\beta 2, \alpha 1, \beta 1$ | $\alpha 1\beta 2\beta 1\alpha 2\alpha 2$ | CC | 0 | WdT1 | 1 | 0.775338 |
| 3 | $\alpha 2\text{-}\alpha 2\text{-}\beta 2, \alpha 1, \beta 1$ | $\alpha 1\alpha 2\beta 1\alpha 2\beta 2$ | WdT1 | 2 | WdT1 | 2 | 0.774371 |
| 4 | $\alpha 2\text{-}\alpha 2\text{-}\beta 2, \alpha 1, \beta 1$ | $\alpha 1\beta 1\alpha 2\alpha 2\beta 2$ | CC | 0 | CC | 0 | 0.759785 |
| 5 | $\alpha 2\text{-}\alpha 2\text{-}\beta 2, \alpha 1, \beta 1$ | $\alpha 1\alpha 2\alpha 2\beta 1\beta 2$ | CC | 0 | WdT1 | 1 | 0.752578 |
| 6 | $\alpha 2\text{-}\alpha 2\text{-}\beta 2, \alpha 1, \beta 1$ | $\alpha 1\beta 2\beta 1\alpha 2\alpha 2$ | CC | 0 | WdT1 | 1 | 0.747161 |

The frequency with which identical subunit orders were predicted from free subunits alone and from linked concatemers was assessed for statistical correlation. A Pearson correlation test revealed significant correlation between the two conditions (Pearson’s correlation coefficient r = 0.7503, p < 0.0001). This analysis included all functional concatemer combinations.

To evaluate structural outcomes of the linked subunit constructs, pentameric AlphaFold2 models were analysed for variations in concatemer orientation and wedge-type formations. Across certain constructs, wedged arrangements were more frequently observed than adjacent arrangements. This was especially noticeable for construct combination α3-α3, β1-β2, β2, where only one out of 15 generated models appeared with both linkers in a counterclockwise, adjacent arrangement. Approximately 21% of the models were predicted to be fully adjacent assemblies. 105 models were predicted to have at least one linker as a *Wedge Type 1,* while *Wedge Type 2* was observed for only for 30 models. Interestingly, the concatemer α2-α2-β2 with the free subunits α1 and β1 had five of 15 models in a counterclockwise adjacent orientation, one of these being the highest scoring model for this arrangement. An overview of this data can be found at https://gitlab.com/hansahls/nachr-stoichiometry-repo/-/blob/main/R_scripts/Linker_Analysis.html as well as in Supplementary Materials I.

## Discussion

This study successfully modelled the ligand-gated ion channels Lsa-nAChR1 and Lsa-nAChR2 from *Lepeophtheirus salmonis* using AlphaFold2 and identified high-scoring stoichiometries with additional use of two protein-protein interaction tools. While bioinformatic predictions provided consistent top-ranking subunit arrangements, experimental results diverged and even conflicted in key cases where constructs used concatenated subunits. AlphaFold2 modelling of these linked constructs revealed conformational variability in the assemblies, such as wedging phenomena. These results challenge the assumption that linker design reliably enforces subunit order, an approach widely used in receptor biology^15,17,22,39–41^.

The discovery of wedged arrangements has implications that extend beyond arthropod nAChRs. Although the findings presented here are based on computational modelling, the consistent prediction of wedged configurations suggests that linker flexibility may permit unexpected subunit arrangements. Given the prevalence of concatemer constructs in receptor research, particularly in GABA_A_, glycine, and nicotinic acetylcholine receptor systems, these results suggest that structural flexibility may be a general phenomenon. Although further experimental validation is required, the structural variability observed in this study raises important questions about the interpretation of concatemer-based studies across receptor systems, in vertebrate and invertebrate systems alike.

Based on the combined evidence from structural modelling, electrophysiological experiments, and sequence alignment, the most likely stoichiometry for Lsa-nAChR1 is α1β1α2α2β2 (Figure 5). This configuration ranked highest in the composite scoring system combining AlphaFold2, PRODIGY, and ZRANK, and achieved the top total robust score. Electrophysiological experiments using a brute-force concatemer approach independently indicated this conformation. Functional receptor expression was observed only when the injected concatemer combination included a linked α2-α2 dimer. Predictive modelling of this linked dimer, in combination with free subunits, produced 12 out of 30 models without wedging and generally higher AlphaFold2 scores of the α2-α2 dimer (Supplementary File I).

**Figure 5:**
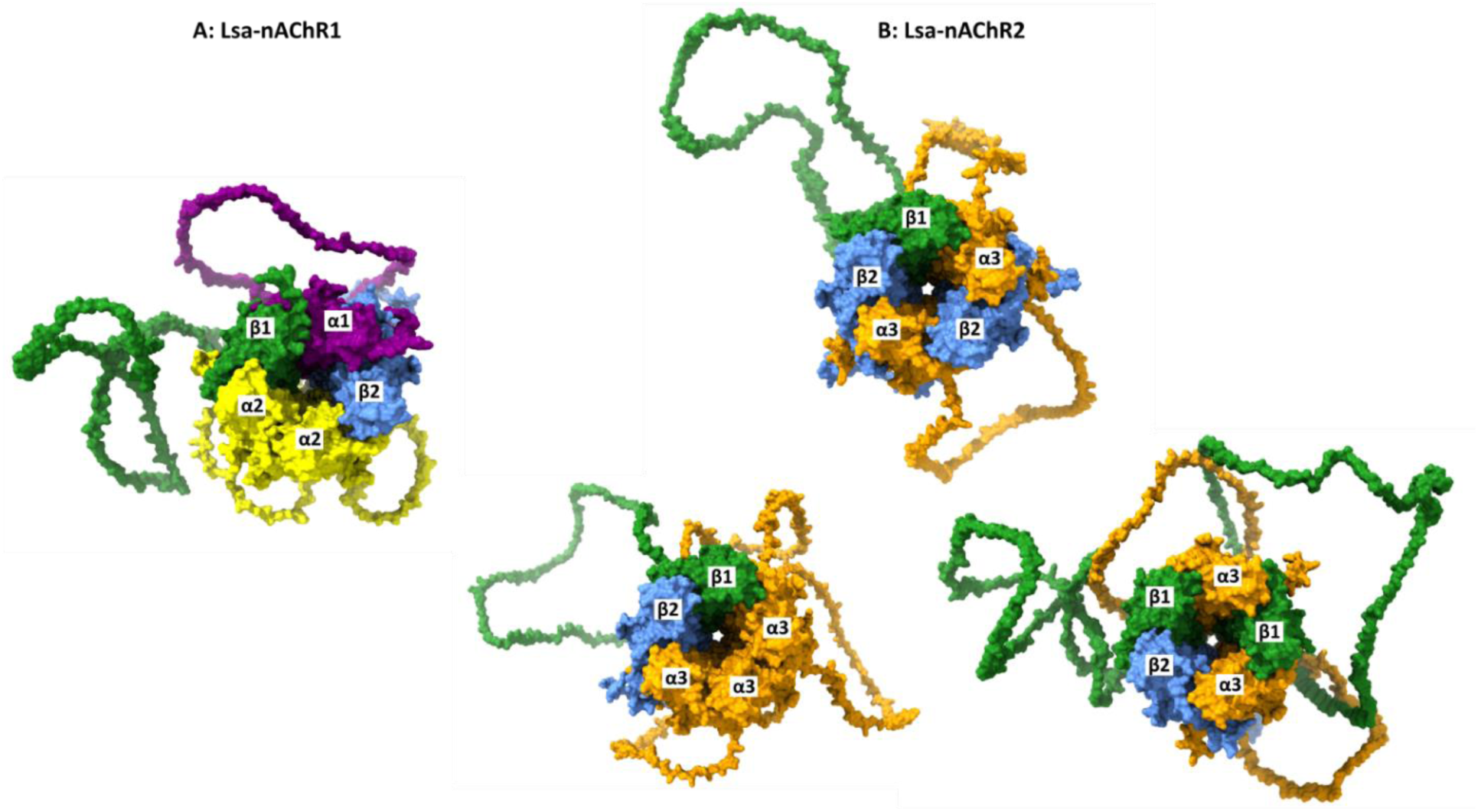
Predicted stoichiometries for Lsa-nAChR1 (A) and Lsa-nAChR2 (B) based on integrated computational and experimental analyses. All receptor models are shown from a top-down, extracellular perspective. In this orientation, the subunits are labelled in a counterclockwise order. The predicted stoichiometry for Lsa-nAChR1 is α1β1α2α2β2, while Lsa-nAChR2 has three functional stoichiometries: α3β1β2α3β2, α3β1α3β1β2, and α3α3α3β1β2. Subunits are color-coded by type (α1 = purple, α2= yellow, α3= orange, β1 = green, β2 = blue), and models were rendered in ChimeraX based on AlphaFold2 structural predictions.

In contrast to Lsa-nAChR1, structural modelling and electrophysiological testing of Lsa-nAChR2 suggest the presence of several functional stoichiometries. Among the AlphaFold2-generated pentamer models, three configurations were identified as strong candidates: α3β1β2α3β2, α3β1β2α3β1 and α3α3α3β1β2. All of these models scored high in the ranking and total robust scaling analysis, reflecting both favourable inter-subunit interfaces and stable overall assembly (Figure 1). The electrophysiological experiments validated three unique compositions, which aligned with the AlphaFold2 predictions. Electrophysiological signals were recorded in oocytes that had been injected with the concatemer combination α3-α3, β1-β2, and β2, forcing the formation of a β2-rich pentamer. Separately, signals were also recorded for the concatemer combinations α3-α3, β1-β1, β2, and β1-β1, α3-β2, α3, forcing the formation of a β1-rich pentamer. Finally, a small signal was recorded for the concatemer β1-β2 when combined with the free subunit α3, forcing the formation of an α3-rich pentamer. These validation experiments demonstrated that the signal obtained when Lsa-nAChR2 was first described in 2020^10^, may be a pooled signal of at least two, possibly three, distinct subunit compositions. At least two distinct subtypes of Lsa-nAChR2 were also suggested by Rufener et al. (2020)^10^.

All three configurations appear structurally feasible; however, it is worth noting that α3β1β2α3β2 achieved the highest combined score from AlphaFold2, PRODIGY, and ZRANK, as well as the highest total robust score. Moreover, it exhibited the highest median total robust score across all models as well as having the most models form in this stoichiometry. When linked concatemers were modelled in AlphaFold2, this arrangement was frequently observed.

However, approximately half of the models displayed wedged features to achieve this stoichiometry, rendering the linker useless as a guide to subunit order. (For all modelled linked concatemers, https://gitlab.com/hansahls/nachr-stoichiometry-repo/-/tree/main/Output_Alphafold2_Computer/Linker_Models).

It is generally assumed that nAChRs need two binding sites for ACh, located in the interface between a α-subunit and an adjacent subunit^4^. Two binding sites are achieved with the α3β1β2α3β2 stoichiometry, at the α3β2 and α3β1 interfaces. The β1-rich stoichiometry α3β1β2α3β1 assembles two identical binding sites, both at the α3β1 interfaces. In the case of the α3-rich pentamer, α3α3α3β1β2, two binding sites can only be obtained by one of them being in the interface between two α3-subunits. Based on the results obtained in 2020^10^, it is unlikely that α3 alone can form a functional channel. The response of the α3-rich composition in the current study was substantially lower compared to the other two compositions. Thus, this configuration may not occur naturally in salmon lice, or is not as preferable as other configurations.

The experimental results for Lsa-nAChR2 composed of different concatenated subunits demonstrated discrepancies. Several concatemer combinations yielded functional receptors, including the trimer β1-β2-α3 when co-expressed with the free subunits α3 and β1. However, when this trimer was combined with linked dimers α3-β1 or β1-α3, no functional receptor was observed. All individual components had previously supported receptor formation when tested independently. One possible explanation is that certain combinations may promote wedged assemblies, as observed in AlphaFold2 models of linked constructs, where wedged formations were more frequently predicted than adjacent formations. Such structural variations may interfere with proper channel assembly or function. Another possible explanation is that the *Xenopus* oocytes failed to express sufficient quantities of both the linked trimer and dimer to achieve a functional receptor on the surface. These findings underscore the limitations of concatemer-based approaches and highlight the need for cautious interpretation of negative functional results, as they may reflect structural interference rather than true incompatibility.

A previously undocumented phenomenon was observed when modelling pentamers with linked dimers and trimers. Predictive modelling indicated that subunits could wedge between a linked dimer pair, defying the intended conformation of the pentamer. These results may cast doubt on previously determined stoichiometries based on concatemer studies^17–22,40,42^. Furthermore, the possibility that some “non-functional” concatemer constructs may be structurally blocked or disrupted by linker interference with the channel pore suggests careful reconsideration. These findings resonate with earlier concerns raised in the broader literature on protein receptors. For example, Liao. et al. ^43^ demonstrated that GABA_A_ receptor concatemers could form as clockwise assemblies, leading them to testing shortening linker lengths to enforce orientation. Earlier reviews and studies have also reported cases of “dangling” subunits, where one subunit of a linked pair incorporates into the pentamer while the other subunit remains excluded^41,44,45^. Although previous studies have highlighted the challenges of clockwise configurations, the specific phenomenon of subunits wedging between linked dimers observed in this study does not appear to have been previously reported. Addressing these limitations through dynamic modelling and cryo-electron microscopy would further refine our understanding of nAChR stoichiometry and guide the interpretation of linker flexibility in previous research on nAChRs and other receptors.

This multifaceted approach for determining nAChR stoichiometry has many strengths, including coherence between *in silico* model scores and recordings from *ex vivo* expression. Nevertheless, some methodological constraints remain. AlphaFold2 achieves high structural fidelity, with a median Global Distance Test Total Score (GDT-TS) > 90 on many CASP14 targets. Furthermore, DeepMind has released a pre-print where AlphaFold-Multimer has demonstrated near-native accuracy for stable complexes, particularly homodimers and conserved heterodimers, with DockQ scores above 0.8^23^. However, its performance declines with larger assemblies involving more than four chains, which is important to consider when interpreting the pentameric models presented here^46^, although this has been improved in recent analyses^47^. Furthermore, AlphaFold generates static models, and does not account for protein dynamics, such as conformational changes in the C-loop during ligand binding, or further cascade conformational changes that open and close the channel pore. Consequently, docking simulations based on these rigid models may not fully capture the energetics of ligand-receptor interactions.

It is worth noting that when AlphaFold-Multimer is used to model linker dimers and trimers as single-chain constructs, wedged conformations are frequently predicted. These findings suggest that these phenomena are not only theoretically plausible but may be common, warranting further investigation in experimental research. Structural techniques such as protein crystallography, cryo-EM, and cross-linking remain essential to verify *in silico* models, providing insight into subunit arrangement, concatemer assembly, and the reliability of AlphaFold predictions.

To characterize the possible binding pockets for Ach in Lsa-nAChR1 and Lsa-nAChR2, multiple sequence alignment comparing vertebrate and invertebrate species was performed. The multiple sequence alignment focused on conserved loop regions A-F, which are known to contribute to ACh binding in nAChRs ^36^. It was hypothesized that *L. salmonis* α1, α2, and α3 would conserve loops A-C, and β1 and β2 would conserve loops D-F^36^. However, the alignment revealed several deviations from this assumption. Most notably, the Lsa-β1 subunit lacked identifiable conservation across all complementary loops, suggesting a limited role in ligand-binding. In contrast, Lsa-β2 retained conserved residues in Loops D and F. Surprisingly, the α1, α2, and α3 subunits also showed conservation in loops D and F. As these are complementary component loops, this raises the possibility that these α-subunits may also contribute to the complementary binding interfaces. This is particularly intriguing for α2 and α3, which, despite possessing both principal and complementary loop features, do not form functional homomeric receptors in the model organisms *D. melanogaster* ^48^ and *L. salmonis*^10^, whereas human α7 is able to function homomerically ^49–51^ without conserved complementary loops. Results presented here show that the configuration α3α3α3β1β2 gives a small response to ACh, indicating that acetylcholine can bind within an α3-α3 binding pocket, or potentially that two binding pockets for ACh is not essential to elicit a small response. This finding supports the idea that β-subunits contribute essential stability or regulatory interactions that enable functional receptor formation in *L. salmonis*.

## Conclusion

In conclusion, this study provides a multidisciplinary approach for predicting and validating the stoichiometry of heteromeric nAChRs from *Lepeophtheirus salmonis*. This was done by integrating AlphaFold2-based structural modelling, protein-protein interaction metrics, sequence conservation analysis, and electrophysiological validation using concatenated subunits. The most likely subunit arrangements for Lsa-nAChR1 and Lsa-nAChR2 were identified as α1β1α2α2β2 for Lsa-nAChR1, and α3β1β2α3β2, α3β1β2α3β1, and α3α3α3β1β2 for Lsa-nAChR2. In addition, these findings reveal phenomena described here as “wedging”, challenging the assumption that linker design alone enforces subunit order and highlight the need for caution when interpreting concatemer-based functional assay results. More broadly, this work underscores the value of combining computational and experimental approaches to resolve complex receptor conformations and opens new avenues for exploring subunit-specific contributions to receptor function.

## Supporting information

All supplementary tables and information provided

## Author Contributions

H.M.S., T.E.H., and K.H.H. performed computational analysis and analysed the data. L.R. contributed to experimental design and setup, and A.S. performed electrophysiological experiments. H.M.S. and

T.E.H. wrote the manuscript. T.E.H. and M.J.B. supervised the project and contributed to manuscript editing. All authors discussed the results and contributed to the final manuscript.

## Competing Interests

None

### Acknowledgement

This study was funded by The Research Council of Norway, project “Development of fully functional, species-specific nicotinic acetylcholine receptor models from arthropods” grant number 325190, and the Norwegian University of Life Sciences.

## Endnote Reference List

1. Gotti, C., Zoli, M. & Clementi, F. Brain nicotinic acetylcholine receptors: native subtypes and their relevance. Trends Pharmacol Sci 27, 482–91 (2006).

2. Arneric, S.P., Holladay, M. & Williams, M. Neuronal nicotinic receptors: a perspective on two decades of drug discovery research. Biochem Pharmacol 74, 1092–101 (2007).

3. Arias, H.R. Localization of agonist and competitive antagonist binding sites on nicotinic acetylcholine receptors. Neurochem Int 36, 595–645 (2000).

4. Millar, N.S. Assembly and subunit diversity of nicotinic acetylcholine receptors. Biochem Soc Trans 31, 869–74 (2003).

5. Costello, M.J. Ecology of sea lice parasitic on farmed and wild fish. Trends Parasitol 22, 475–83 (2006).

6. Costello, M.J. The global economic cost of sea lice to the salmonid farming industry. J Fish Dis 32, 115–8 (2009).

7. Torrissen, O. et al. Salmon lice – impact on wild salmonids and salmon aquaculture. Journal of Fish Diseases 36, 171–194 (2013).

8. Wagner, C.A., Friedrich, B., Setiawan, I., Lang, F. & Bröer, S. The Use of Xenopus laevis Oocytes for the Functional Characterization of Heterologously Expressed Membrane Proteins. Cell Physiol Biochem 10, 1–12 (2000).

9. Matsuda, K., Ihara, M. & Sattelle, D.B. Neonicotinoid Insecticides: Molecular Targets, Resistance, and Toxicity. Annu Rev Pharmacol Toxicol 60, 241–255 (2020).

10. Rufener, L., Kaur, K., Sarr, A., Aaen, S.M. & Horsberg, T.E. Nicotinic acetylcholine receptors: Ex-vivo expression of functional, non-hybrid, heteropentameric receptors from a marine arthropod, Lepeophtheirus salmonis. PLoS Pathog 16, e1008715 (2020).

11. Chang, Y., Wang, R., Barot, S. & Weiss, D.S. Stoichiometry of a recombinant GABAA receptor. J Neurosci 16, 5415–24 (1996).

12. Farrar, S.J., Whiting, P.J., Bonnert, T.P. & McKernan, R.M. Stoichiometry of a ligand-gated ion channel determined by fluorescence energy transfer. J Biol Chem 274, 10100–4 (1999).

13. Drenan, R.M. et al. Subcellular trafficking, pentameric assembly, and subunit stoichiometry of neuronal nicotinic acetylcholine receptors containing fluorescently labeled alpha6 and beta3 subunits. Mol Pharmacol 73, 27–41 (2008).

14. Durisic, N. et al. Stoichiometry of the human glycine receptor revealed by direct subunit counting. J Neurosci 32, 12915–20 (2012).

15. Baumann, S.W., Baur, R. & Sigel, E. Subunit arrangement of gamma-aminobutyric acid type A receptors. J Biol Chem 276, 36275–80 (2001).

16. Groot-Kormelink, P.J., Broadbent, S., Beato, M. & Sivilotti, L.G. Constraining the expression of nicotinic acetylcholine receptors by using pentameric constructs. Mol Pharmacol 69, 558–63 (2006).

17. Baumann, S.W., Baur, R. & Sigel, E. Forced Subunit Assembly in α1β2γ2 GABAAReceptors: INSIGHT INTO THE ABSOLUTE ARRANGEMENT*. Journal of Biological Chemistry 277, 46020–46025 (2002).

18. Kaur, K.H., Baur, R. & Sigel, E. Unanticipated structural and functional properties of delta-subunit-containing GABAA receptors. J Biol Chem 284, 7889–96 (2009).

19. Botzolakis, E.J. et al. Comparison of γ-Aminobutyric Acid, Type A (GABAA), Receptor αβγ and αβδ Expression Using Flow Cytometry and Electrophysiology: EVIDENCE FOR ALTERNATIVE SUBUNIT STOICHIOMETRIES AND ARRANGEMENTS*. Journal of Biological Chemistry 291, 20440–20461 (2016).

20. Im, W.B., Pregenzer, J.F., Binder, J.A., Dillon, G.H. & Alberts, G.L. Chloride Channel Expression with the Tandem Construct of α6-β2 GABAA Receptor Subunit Requires a Monomeric Subunit of α6 or γ2 (∗). The Journal of biological chemistry 270, 26063–26066 (1995).

21. Bracamontes, J.R. & Steinbach, J.H. Steroid Interaction with a Single Potentiating Site Is Sufficient to Modulate GABA-A Receptor Function. Mol Pharmacol 75, 973–981 (2009).

22. Shu, H.-J. et al. Characteristics of concatemeric GABAA receptors containing α4/δ subunits expressed in Xenopus oocytes. British Journal of Pharmacology 165, 2228–2243 (2012).

23. Evans, R., et al. Protein complex prediction with AlphaFold-Multimer. *bioRxiv* (2022).

24. Jumper, J. et al. Applying and improving AlphaFold at CASP14. Proteins 89, 1711–1721 (2021).

25. Jumper, J. et al. Highly accurate protein structure prediction with AlphaFold. Nature 596, 583–589 (2021).

26. Jeppe Hallgren, K.D.T., Mads D. Pedersen, José Juan Almagro Armenteros, Paolo Marcatili, Henrik Nielsen, Anders Krogh and Ole Winther. DeepTMHMM predicts alpha and beta transmembrane proteins using deep neural networks. Journal of Engineering, 401 (2022).

27. Pierce, B. & Weng, Z. ZRANK: reranking protein docking predictions with an optimized energy function. Proteins 67, 1078–86 (2007).

28. Vangone, A. & Bonvin, A. PRODIGY: A Contact-based Predictor of Binding Affinity in Protein-protein Complexes. Bio Protoc 7, e2124 (2017).

29. Qian, H.J. et al. RobustScaler: QoS-Aware Autoscaling for Complex Workloads. 2022 Ieee 38th International Conference on Data Engineering (Icde 2022), 2762–2775 (2022).

30. Staudte, R.G. Inference for the Standardized Median. in Contemporary Developments in Statistical Theory 353–363 (2014).

31. Moradi, E., Elsisi, M., Mahmoud, K., Lehtonen, M. & Darwish, M.M.F. Robust deep neural network-based internet of things for power transformer fault diagnosis under imbalanced data and uncertainties. International Journal of Electrical Power & Energy Systems 168(2025).

32. Ordoñez-Avila, R., Meza, J. & Ventura, S. Mining autonomous student patterns score on LMS within online higher education. Peerj Computer Science 11(2025).

33. Wongoutong, C. The impact of neglecting feature scaling in k-means clustering. Plos One 19(2024).

34. Pettersen, E.F. et al. UCSF Chimera--a visualization system for exploratory research and analysis. J Comput Chem 25, 1605–12 (2004).

35. Okonechnikov, K., Golosova, O., Fursov, M. & Team, U. Unipro UGENE: a unified bioinformatics toolkit. Bioinformatics 28, 1166–1167 (2012).

36. Grutter, T. & Changeux, J.-P. Nicotinic receptors in wonderland. Trends in Biochemical Sciences 26, 459–463 (2001).

37. Genz, L.R., Nair, S., Nagar, N. & Topf, M. Assessing scoring metrics for AlphaFold2 and AlphaFold3 protein complex predictions. bioRxiv (2025).

38. Homma, F., Huang, J. & van der Hoorn, R.A.L. AlphaFold-Multimer predicts cross-kingdom interactions at the plant-pathogen interface. Nat Commun 14, 6040 (2023).

39. White, M.M. Pretty Subunits all in a row: Using concatenated subunit constructs to force the expression of receptors with defined subunit stoichiometry and spatial arrangement. Molecular Pharmacology 69, 407–410 (2006).

40. Minier, F. & Sigel, E. Techniques: Use of concatenated subunits for the study of ligand-gated ion channels. Trends Pharmacol Sci 25, 499–503 (2004).

41. Zhou, Y. et al. Human alpha4beta2 acetylcholine receptors formed from linked subunits. J Neurosci 23, 9004–15 (2003).

42. Baumann, S.W., Baur, R. & Sigel, E. Individual properties of the two functional agonist sites in GABA(A) receptors. J Neurosci 23, 11158–66 (2003).

43. Liao, V.W.Y. et al. Concatenated γ-aminobutyric acid type A receptors revisited: Finding order in chaos. Journal of General Physiology 151, 798–819 (2019).

44. J. Boileau, S.S.E.a.A. Tandem Couture: Cys-loop receptor concatemer insights and caveats. Molecular Neurobiology, 113–128 (2007).

45. Groot-Kormelink, P.J., Broadbent, S.D., Boorman, J.P. & Sivilotti, L.G. Incomplete incorporation of tandem subunits in recombinant neuronal nicotinic receptors. J Gen Physiol 123, 697–708 (2004).

46. Zhu, W., Shenoy, A., Kundrotas, P. & Elofsson, A. Evaluation of AlphaFold-Multimer prediction on multi-chain protein complexes. Bioinformatics 39(2023).

47. Bryant, P. & Noe, F. Improved protein complex prediction with AlphaFold-multimer by denoising the MSA profile. PLoS Comput Biol 20, e1012253 (2024).

48. Komori, Y. et al. Functional impact of subunit composition and compensation on Drosophila melanogaster nicotinic receptors-targets of neonicotinoids. PLoS Genet 19, e1010522 (2023).

49. Conroy, W.G.B., Darwin K. Neurons Can Maintain Multiple Classes of Nicotinic Acetylcholine Receptors Distinguished by Different Subunit Compositions (∗). The Journal of biological chemistry 270, 4424–4431 (1995).

50. Kalamida, D. et al. Muscle and neuronal nicotinic acetylcholine receptors. Structure, function and pathogenicity. FEBS J 274, 3799–845 (2007).

51. Lindstrom, J.M. Acetylcholine receptors and myasthenia. Muscle & Nerve 23, 453–477 (2000).

