## Supplementary material for "AlphaFold-Multimer Modelling of Linked nAChR Subunits Challenges Concatemer Design Assumptions": All supplementary tables and information provided

### Supplementary Materials

#### A: Gitlab Repository:

https://gitlab.com/hansahls/nachr-stoichiometry-repo

#### B: Gene IDs and RefSeqs of Lsa-nAChR subunits

Table 1

Gene IDs and RefSeqs for each of the subunits included in this study.

| Gene ID | RefSeq (mRNA Transcript) | Protein | Description | Source |
|---|---|---|---|---|
| 121121295 | XM_040716185.2 | XP_040572119.1 | nAChRa1 | NCBI RefSeq/ GenBank |
| 121121811 | XM_040716792.2 | 040572726.1 | nAChRa2 | NCBI RefSeq/ GenBank |
| 121121555 | XM_040716513.2 | XP_040572447.1 | nAChRa3 | NCBI RefSeq/ GenBank |
| 121115977 | XM_040710162.2 | XP_040566096.1 | nAChRb1 | NCBI RefSeq/ GenBank |
| 121115804 | XM_040709951.2 | XP_040565885.1 | nAChRb2 | NCBI RefSeq/ GenBank |

#### C: Primer design of linked subunits

Table 2

Primers used in constructing the plasmids for injection into X. laevis. The primer name and sequence are specified, along with whether the primer was used to construct a monomer, dimer, or trimer.

| Primer name | Sequence 5' to 3' | Remark |
|---|---|---|
| NheI_Ls-nAChRa1F1 | GGCGGCTAGCAGATTTCGCATCCTCTCCAA | Monomer |
| XhoI_Ls-nAChRa1R1 | GGCGCTCGAGTTTTGGCTTCTTCTTCTTCTTCA | Monomer |
| NheI_Ls-nAChRb1F1 | GGCGGCTAGCCATCATCAAGAATGGATTGGAA | Monomer |
| XhoI_Ls-nAChRb1R1 | GGCGCTCGAGTGCAGCTGTATTTCCTTCTTCT | Monomer |
| NheI_Ls_nAChRa3F1 | GGCGGCTAGCGAAAACATGGACAAAGTTTGGA | Monomer |
| Xhol_Ls_nAChRa3R1 | GGCGCTCGAGGGAGGGTGGGGTGTAGGTAT | Monomer |
| NheI_Ls_nAChRa2F1 | GGCGGCTAGCAAGAAGGGATCGAAAATGCTT | Monomer |
| Xhol_Ls_nAChRa2R1 | GGCGCTCGAGTGAGCAAGAGGATGTTTTCATT | Monomer |
| NheI_Ls_nAChRb2F1 | GGCGGCTAGCGCATAGCGTTTCAAATGTTTCTT | Monomer |
| Xhol_Ls_nAChRb2R1 | GGCGCTCGAGGCAAATGGGTGGGATGATAC | Monomer |
| Ls-a1_linker_R1 | CGCCCTCGAGGGGCCCACTTCCGGCTCGTTGACTTTTAAGTGATTCAGG | Dimers |
| Ls-a1_woSP_F1 | GGCGGCTAGCACCGGTCCTAGGCTTAAGGGGCCCAATCCAGATGCCAAGAGATTAT | Dimers |
| Ls-a2_linker_R1 | CGCCCTCGAGGGGCCCACTTCCGGCGAAAACTTTTTCTTGAGATACTCTG | Dimers |
| Ls-a2_woSP_F1 | GGCGGCTAGCACCGGTCCTAGGCTTAAGGGGCCCAATCCGGATGCAAAGAGACT | Dimers |
| Ls-a3_linker_R1 | CGCCCTCGAGGGGCCCTCCACTTCCGGCTTTCAGGAATGAAAATTTGTTT | Dimers |
| Ls-a3_woSP_F1 | GGCGGCTAGCACCGGTCCTAGGCTTAAGGGGCCCAATCCAGACGCCAAACGACT | Dimers |
| Ls-b1_linker_R1 | CGCCCTCGAGGGGCCCACTTCCGGCACTTCCGGCACTTCCGGCACTTCCGGCCTTTCCTTTATAAATGTCTA TGATCT | Dimers |
| Ls-b1_woSP_F1 | GGCGGCTAGCACCGGTCCTAGGCTTAAGGGGCCCTCACAAGAGGAGGAAAGACTCG | Dimers |

| | | |
|---|---|---|
| Ls-b2_linker_R1 | CGCCCTCGAGGGGCCCACTTCCGGCACTTCCGGCACTTCCGGCACTTCCGGCTTTCGGAATCTCGGAAAGC | Dimers |
| Ls-b2_woSP_F1 | GGCGGCTAGCACCGGTCCTAGGCTTAAGGGGCCCAATCCTCATGCCAAACGACT | Dimers |
| Ls-a1_linker_AflII | CGCCCTCGAGCTTAAGGGCTCGTTGACTTTTAAGTGATTCAGG | Dimers + add AflII to make trimers |
| Ls-a2_linker_AflII | CGCCCTCGAGCTTAAGGGCGAAAACTTTTTCTTGAGATACTCTG | Dimers + add AflII to make trimers |
| Ls-a3_linker_AflII | CGCCCTCGAGCTTAAGTCCGGCTTTCAGGAATGAAAATTTGTTT | Dimers + add AflII to make trimers |
| Ls-b1_linker_AflII | CGCCCTCGAGCTTAAGGGCACTTCCGGCACTTCCGGCACTTCCGGCCTTTCCTTTATAAATGTCTATGATCT | Dimers + add AflII to make trimers |
| Ls-b2_linker_AflII | CGCCCTCGAGCTTAAGGGCACTTCCGGCACTTCCGGCACTTCCGGCTTTCGGAATCTCGGAAAGC | Dimers + add AflII to make trimers |
| Lsa-a1_casette-F1 | GGCGACCGGTGCCGGAAGTGCCGGAAGTAATCCAGATGCCAAGAGA | Monomer without START and STOP for trimers |
| Ls-a1_casette-R1 | CGCCGGGCCCACTTCCGGCACTTCCGGCACTTCCGGCACTTCCGGCTCGTTGACTTTTAAGTGATTCAGG | Monomer without START and STOP for trimers |
| Lsa-b2_casette-F1 | GGCGACCGGTGCCGGAAGTGCCGGAAGTAATCCTCATGCCAAACGACT | Monomer without START and STOP for trimers |
| Ls-b2_casette-R1 | CGCCGGGCCCACTTCCGGCACTTCCGGCACTTCCGGCACTTCCGGCACTTCCGGCACTTCCGGCTTTCGG | Monomer without START and STOP for trimers |
| Lsa-a2_casette-F1 | GGCGACCGGTGCCGGAAGTGCCGGAAGTAATCCGGATGCAAAGAGACT | Monomer without START and STOP for trimers |
| Ls-a2_casette-R1 | GGGCCCACTTCCGGCACTTCCGGCACTTCCGGCACTTCCGGCGAAAACTTTTTCTTGAGATACTCTG | Monomer without START and STOP for trimers |

### D: Linker sequences (including parts of the C- and N-terminus)

Linker a3_a3: AGSGGP

Linker b1_b2: AGSAGSAGSAGSPGTRPTTGPRLKGP

Linker b1_b1: AGSAGSAGSAGSPGTRPTTGPRLKGP

Linker a3_b2: AGSGGP

Linker a2_a2_b2.1: VSPIDVIFSKIALEESSRVSQEKVFPGTRPTTGAGSAGS

Linker a2_a2_b2.2: AGSAGSAGSGP

Linker b1_b2_a3.1: AGSAGSAGSAGSPGTRPTTGPRLKGP

Linker b1_b2_a3.2: AGSAGSAGSAGSGP

### E: Rankings of Lsa-nAChR1 Pentamer Models

Table 3

Ranks of all pentamer models containing the subunits α1, α2, β1 and β2 in the nicotinic acetylcholine receptor Lsa-nAChR1 from *Lepeophtheirus salmonis*. The pentamers were generated and scored with AlphaFold2, while individual dimers in the pentamers were scored with ZRANK and PRODIGY and then summed up for each. The Ranksum represents the sum of individual ranks for AlphaFold2, PRODIGY and ZRANK. The stoichiometries are sorted by Rank of Ranksum and presented in a counterclockwise order when viewed from the synapsis. The best scoring model per stoichiometry is indicated in bold red.

| Stoichiometry | AlphaFold2 Score | PRODIGY Score | ZRANK Score | AlphaFold2 Rank | PRODIGY Rank | ZRANK Rank | Ranksum | Rank of Ranksum | Number per Stoichiometry |
|---|---|---|---|---|---|---|---|---|---|
| **α1β1α2α2β2** | **0.765639013** | **-121.3** | **-1883.291** | **12** | **1** | **5** | **18** | **1.5** | 5 |
| **α1α2β2β1β2** | **0.775139217** | **-113.1** | **-1901.115** | **8** | **6** | **4** | **18** | **1.5** | 2 |
| **α1β2α2β1β2** | **0.77587734** | **-109.4** | **-1909.744** | **6** | **9.5** | **3** | **18.5** | **3** | 7 |
| α1α2β2β1β2 | 0.780962955 | -111.5 | -1827.646 | 1 | 8 | 10 | 19 | 4 | |
| α1β1α2α2β2 | 0.764297179 | -113.7 | -1801.721 | 14 | 5 | 13 | 32 | 5 | |
| α1β2α2β1β2 | 0.771459045 | -106.4 | -1852.948 | 9 | 25 | 6 | 40 | 6 | |

| | | | | | | | | | |
|---|---|---|---|---|---|---|---|---|---|
| **α1β1β2β2α2** | **0.752849716** | **-117.3** | **-1809.232** | **28** | **3** | **11** | **42** | **7** | 3 |
| **α1β1β2α2β1** | **0.747960286** | **-118** | **-1957.076** | **43** | **2** | **2** | **47** | **8** | 4 |
| **α1β1α2β1β2** | **0.747735257** | **-114.6** | **-1962.209** | **44** | **4** | **1** | **49** | **9** | 12 |
| **α1α1β1β2α2** | **0.75810183** | **-109.3** | **-1775.927** | **21** | **11.5** | **18** | **50.5** | **10** | 9 |
| **α1α2β2α2β1** | **0.755628906** | **-109.4** | **-1727.366** | **24** | **9.5** | **20** | **53.5** | **11** | 3 |
| α1β1α2β1β2 | 0.752484252 | -108.5 | -1805.383 | 31 | 14 | 12 | 57 | 12.5 | |
| α1β1β2α2β1 | 0.74808022 | -112.4 | -1834.222 | 42 | 7 | 8 | 57 | 12.5 | |
| **α1α2α1β1β2** | **0.752602995** | **-108.5** | **-1797.548** | **29** | **14** | **15** | **58** | **14** | 4 |
| **α1β1α1β2α2** | **0.751612087** | **-107.8** | **-1830.108** | **33** | **16.5** | **9** | **58.5** | **15** | 3 |
| **α1α2β1β2β2** | **0.769252527** | **-104.9** | **-1778.763** | **10** | **37.5** | **17** | **64.5** | **16** | 8 |
| **α1β1β2α2β2** | **0.777839625** | **-104.6** | **-1718.957** | **4** | **40** | **21** | **65** | **17** | 1 |
| **α1α2β1β2β1** | **0.7459293** | **-109.3** | **-1834.595** | **49** | **11.5** | **7** | **67.5** | **18** | 2 |
| α1α2β2α2β1 | 0.75873419 | -107.1 | -1676.805 | 20 | 21.5 | 27 | 68.5 | 19 | |
| α1α2β2α2β1 | 0.762998345 | -106.5 | -1658.883 | 15 | 24 | 33 | 72 | 20 | |
| **α1α2α2β1β2** | **0.761684675** | **-107.2** | **-1637.728** | **17** | **19** | **39** | **75** | **21** | 5 |
| α1β2α2β1β2 | 0.780512493 | -101.8 | -1716.634 | 2 | 58.5 | 22 | 82.5 | 22 | |
| α1α2α1β1β2 | 0.753756071 | -104.3 | -1800.71 | 27 | 42 | 14 | 83 | 23 | |
| **α1α2α1β2β1** | **0.751014037** | **-105** | **-1780.44** | **34** | **34** | **16** | **84** | **24** | 2 |
| **α1β1α2β2β2** | **0.757940716** | **-105.8** | **-1654.286** | **22** | **27** | **36** | **85** | **25** | 3 |
| **α1α2β2β1α2** | **0.775196185** | **-108.5** | **-1570.407** | **7** | **14** | **65** | **86** | **26** | 1 |
| α1α2β1β2β2 | 0.778096736 | -101.7 | -1709.089 | 3 | 61 | 23 | 87 | 27 | |
| α1β2α2β1β2 | 0.777736457 | -101.8 | -1672.122 | 5 | 58.5 | 29 | 92.5 | 28 | |
| α1α2β1β2β2 | 0.768422269 | -100.4 | -1661.401 | 11 | 70 | 32 | 113 | 29 | |
| **α1β1β2α2α2** | **0.759245436** | **-105.5** | **-1566.364** | **19** | **30.5** | **67** | **116.5** | **30** | 5 |
| α1β1α2β1β2 | 0.749510645 | -102.1 | -1689.746 | 37 | 56 | 24 | 117 | 31 | |
| α1β1α2α2β2 | 0.748778996 | -106.2 | -1572.683 | 39 | 26 | 61 | 126 | 32 | |
| **α1β1β1β2α2** | **0.722181542** | **-107.8** | **-1747.343** | **91** | **16.5** | **19** | **126.5** | **33** | 3 |
| α1β1α2β2β2 | 0.751995641 | -103.8 | -1591.299 | 32 | 44 | 52 | 128 | 34 | |
| α1β1β2β2α2 | 0.748779757 | -103.2 | -1618.396 | 38 | 48 | 45 | 131 | 35 | |
| α1β1α2α2β2 | 0.743390399 | -107.2 | -1573.099 | 53 | 20 | 60 | 133 | 36 | |
| α1β1β2α2α2 | 0.738701935 | -107.1 | -1580.883 | 60 | 21.5 | 55 | 136.5 | 37 | |
| α1β1β2α2α2 | 0.738645563 | -106.9 | -1582.843 | 61 | 23 | 54 | 138 | 38 | |
| α1α2β1β2β2 | 0.746774333 | -102 | -1640.768 | 45 | 57 | 38 | 140 | 39 | |
| α1α2β1β2β1 | 0.720535344 | -107.3 | -1665.991 | 93 | 18 | 31 | 142 | 40 | |
| **α1α2β2α1β1** | **0.729235666** | **-104.9** | **-1675.926** | **77** | **37.5** | **28** | **142.5** | **41** | 1 |
| α1β1β2α2β1 | 0.724821486 | -105.5 | -1685.942 | 87 | 32 | 25 | 144 | 42 | |
| α1α1β1β2α2 | 0.734905861 | -104.1 | -1657.804 | 69 | 43 | 34 | 146 | 43 | |
| α1β1β2α2β1 | 0.746693472 | -102.7 | -1593.732 | 47 | 50 | 51 | 148 | 44 | |
| α1β1α2β1β2 | 0.727036003 | -105.1 | -1655.897 | 82 | 33 | 35 | 150 | 45 | |
| α1β1α2β1β2 | 0.729452025 | -104.7 | -1648.496 | 76 | 39 | 37 | 152 | 46 | |
| α1β1α2β1β2 | 0.727765694 | -105.5 | -1622.767 | 79 | 30.5 | 43 | 152.5 | 47 | |
| α1α2β1β2β2 | 0.739119263 | -102.7 | -1613.845 | 59 | 50 | 46 | 155 | 48.5 | |
| α1α2β1β2β2 | 0.76203086 | -100.8 | -1539.91 | 16 | 66 | 73 | 155 | 48.5 | |
| α1β1α2β1β2 | 0.723344607 | -104.9 | -1671.309 | 90 | 35.5 | 30 | 155.5 | 50 | |
| α1β1α2β1β2 | 0.723847752 | -104.5 | -1680.457 | 89 | 41 | 26 | 156 | 51 | |
| **α1α2β1β2α2** | **0.754951145** | **-103.6** | **-1504.934** | **26** | **45** | **86** | **157** | **52** | 1 |
| α1β1α2β2β2 | 0.750169289 | -101.5 | -1574.752 | 36 | 63 | 59 | 158 | 53.5 | |
| α1β2α2β1β2 | 0.755993338 | -99.7 | -1575.79 | 23 | 77 | 58 | 158 | 53.5 | |
| **α1β2β1β2α2** | **0.750492237** | **-101** | **-1572.234** | **35** | **64** | **62** | **161** | **55** | 1 |
| α1β1β2α2α2 | 0.746126532 | -103.3 | -1558.414 | 48 | 46 | 69 | 163 | 56 | |
| α1β2α2β1β2 | 0.764955536 | -98.3 | -1567.241 | 13 | 89 | 66 | 168 | 57 | |
| **α1α2α2β2β1** | **0.755000186** | **-103.3** | **-1466.67** | **25** | **47** | **97** | **169** | **58** | 1 |
| α1β1α2α2β2 | 0.73789291 | -105.7 | -1532.169 | 64 | 28 | 78 | 170 | 59 | |
| α1β1β2α2α2 | 0.738586804 | -104.9 | -1536.023 | 62 | 35.5 | 74 | 171.5 | 60 | |
| α1β1α2β1β2 | 0.731687504 | -102.7 | -1603.391 | 74 | 52 | 48 | 174 | 61 | |
| α1α2β1β2β2 | 0.744212738 | -99.7 | -1600.527 | 50 | 75.5 | 50 | 175.5 | 62 | |
| α1α1β1β2α2 | 0.730106118 | -102.3 | -1610.122 | 75 | 54 | 47 | 176 | 63 | |

| | | | | | | | | | |
|---|---|---|---|---|---|---|---|---|---|
| α1α2α2β1β2 | 0.736047131 | -105.6 | -1527.575 | 68 | 29 | 80 | 177 | 64 | |
| α1α2α1β1β2 | 0.743678885 | -98.6 | -1620.913 | 52 | 84 | 44 | 180 | 65 | |
| **α1α2β1α1β2** | **0.739659645** | **-99.2** | **-1587.992** | **57** | **79.5** | **53** | **189.5** | **66** | 2 |
| α1α2α2β1β2 | 0.737446956 | -102.7 | -1535.64 | 65 | 50 | 75 | 190 | 67 | |
| α1β1α2β1β2 | 0.724925153 | -100.8 | -1628.41 | 86 | 65 | 42 | 193 | 68 | |
| α1α2α2β1β2 | 0.744168648 | -102.4 | -1493.893 | 51 | 53 | 90 | 194 | 69.5 | |
| α1β1α2β1β2 | 0.718660876 | -101.8 | -1632.805 | 94 | 60 | 40 | 194 | 69.5 | |
| α1α2β1β2β2 | 0.759479146 | -99 | -1477.318 | 18 | 82 | 96 | 196 | 71.5 | |
| α1β1α2β1β2 | 0.725591247 | -101.6 | -1602.165 | 85 | 62 | 49 | 196 | 71.5 | |
| **α1β2β1α2α2** | **0.752526178** | **-99.9** | **-1482.056** | **30** | **74** | **93** | **197** | **73** | 2 |
| α1β1β2β2α2 | 0.748090914 | -99.5 | -1521.723 | 41 | 78 | 83 | 202 | 74 | |
| **α1α2β1β1β2** | **0.702240301** | **-100.2** | **-1630.888** | **97** | **71.5** | **41** | **209.5** | **75** | 3 |
| α1α1β1β2α2 | 0.727741869 | -100.6 | -1571.945 | 80 | 67.5 | 63 | 210.5 | 76 | |
| **α1β2α2α2β1** | **0.737062135** | **-102.3** | **-1486.661** | **66** | **55** | **91** | **212** | **77.5** | 2 |
| α1β2α2β1β2 | 0.748175991 | -98.4 | -1505.747 | 40 | 87 | 85 | 212 | 77.5 | |
| α1α1β1β2α2 | 0.726013917 | -100.2 | -1580.02 | 84 | 73 | 56 | 213 | 79 | |
| α1β1α1β2α2 | 0.74670505 | -94.6 | -1543.681 | 46 | 99 | 71 | 216 | 80 | |
| α1α2α1β1β2 | 0.736084018 | -99.2 | -1535.007 | 67 | 79.5 | 76 | 222.5 | 81 | |
| **α1α1α2β1β2** | **0.72141731** | **-99.7** | **-1576.2** | **92** | **75.5** | **57** | **224.5** | **82.5** | 2 |
| α1α2α2β1β2 | 0.742377863 | -100.2 | -1455.456 | 54 | 71.5 | 99 | 224.5 | 82.5 | |
| **α1β2α2β1β1** | **0.734020283** | **-98.5** | **-1541.184** | **72** | **85** | **72** | **229** | **84** | 1 |
| α1α2β1α1β2 | 0.741010604 | -97.3 | -1525.166 | 55 | 92.5 | 82 | 229.5 | 85 | |
| α1α1α2β1β2 | 0.72758636 | -99.1 | -1547.724 | 81 | 81 | 70 | 232 | 86 | |
| **α1α1β2α2β1** | **0.737948766** | **-95.5** | **-1530.127** | **63** | **96.5** | **79** | **238.5** | **87.5** | 2 |
| α1β2α2α2β1 | 0.734742816 | -100.6 | -1406.577 | 71 | 67.5 | 100 | 238.5 | 87.5 | |
| α1α1β1β2α2 | 0.7283655 | -98.4 | -1534.94 | 78 | 86 | 77 | 241 | 89 | |
| α1α1β1β2α2 | 0.739730998 | -94 | -1482.689 | 56 | 100 | 92 | 248 | 90 | |
| α1α2β1β1β2 | 0.699533281 | -97.8 | -1571.692 | 98 | 90 | 64 | 252 | 91.5 | |
| α1β1β1β2α2 | 0.726996768 | -98.3 | -1525.579 | 83 | 88 | 81 | 252 | 91.5 | |
| α1α2α1β2β1 | 0.739249701 | -95.5 | -1461.358 | 58 | 96.5 | 98 | 252.5 | 93 | |
| α1α2β1β1β2 | 0.71622639 | -97.4 | -1561.02 | 95 | 91 | 68 | 254 | 94 | |
| α1α1β1β2α2 | 0.723938882 | -98.8 | -1509.042 | 88 | 83 | 84 | 255 | 95 | |
| α1β2β1α2α2 | 0.659074056 | -100.5 | -1501.358 | 100 | 69 | 88 | 257 | 96 | |
| α1α1β2α2β1 | 0.734855529 | -96.5 | -1481.058 | 70 | 94 | 94 | 258 | 97 | |
| α1α1β1β2α2 | 0.732069259 | -96.2 | -1479.732 | 73 | 95 | 95 | 263 | 98 | |
| α1β1α1β2α2 | 0.695013733 | -97.3 | -1503.762 | 99 | 92.5 | 87 | 278.5 | 99 | |
| α1β1β1β2α2 | 0.714153804 | -95.4 | -1498.832 | 96 | 98 | 89 | 283 | 100 | |

### F: Rankings of Lsa-nAChR1 Dimer Models

Table 4

Ranks of the best dimers retrieved from complete pentamers, scored with PRODIGY and ZRANK. Ranking is based on the sum of individual scores for PRODIGY and ZRANK. The stoichiometry is presented in a counterclockwise order when viewed from the synapsis.

| DIMER | PRODIGY SCORE | ZRANK SCORE | RANK |
|---|---|---|---|
| α1_α1 | -21.9 | -352.819 | 11 |
| α1_α2 | -21.7 | -315.827 | 15 |
| α1_β1 | -27.2 | -506.022 | 2 |
| α1_β2 | -21 | -415.296 | 10 |

| | | | |
|---|---|---|---|
| α2_α1 | -24.4 | -383.27 | 6 |
| α2_α2 | -24.5 | -311.512 | 14 |
| α2_β1 | -27.9 | -435.533 | 3 |
| α2_β2 | -24.6 | -364.562 | 8 |
| β1_α1 | -22.5 | -383.821 | 9 |
| β1_α2 | -24.9 | -412.329 | 5 |
| β1_β1 | -20.4 | -322.167 | 16 |
| β1_β2 | -35.4 | -532.561 | 1 |
| β2_α1 | -25.8 | -437.866 | 4 |
| β2_α2 | -22.1 | -347.826 | 12 |
| β2_β1 | -24.5 | -379.477 | 7 |
| β2_β2 | -20.8 | -351.783 | 13 |

### G: Rankings of Lsa-nAChR2 Pentamer Models

Table 5

Ranks of the best unique pentamers containing subunits α3, β1, and β2 in the nicotinic acetylcholine receptor R2 from *Lepeoptheirus salmonis*. The pentamers were generated and scored with AlphaFold2, while individual dimers in the pentamers were scored with ZRANK and PRODIGY and then summed up for each. The Ranksum represents the sum of individual ranks for AlphaFold2, PRODIGY and ZRANK. The stoichiometries are sorted by Rank of Ranksum and presented in a counterclockwise order when viewed from the synapsis. The best scoring model per stoichiometry is indicated in bold red.

| Stoichiometry | AlphaFold2 Score | PRODIGY Score | ZRANK Score | AlphaFold2 Rank | ZRANK Rank | PRODIGY Rank | Ranksum | Rank of Ranksum | Number of Stoichiometry |
|---|---|---|---|---|---|---|---|---|---|
| **α3β1β2α3β2** | **0.784236466** | **-105.8** | **-1839.721** | **2** | **7** | **9** | **18** | **1** | 16 |
| α3β1β2α3β2 | 0.783174664 | -106.4 | -1838.221 | 3 | 8 | 7 | 18 | 2 | |
| α3β1β2α3β2 | 0.779879184 | -104.8 | -1808.859 | 6 | 10 | 11 | 27 | 3 | |
| α3β1β2α3β2 | 0.783023057 | -102 | -1821.56 | 4 | 9 | 15 | 28 | 4 | |
| α3β1β2α3β2 | 0.776741476 | -103.8 | -1797.664 | 9 | 11 | 13 | 33 | 5 | |
| α3β1β2α3β2 | 0.757137661 | -106.5 | -1868.454 | 26 | 5 | 6 | 37 | 6 | |
| **α3α3α3β1β2** | **0.754718136** | **-111.8** | **-1919.649** | **35** | **2** | **1** | **38** | **7** | 15 |
| α3β1β2α3β2 | 0.792443854 | -96.5 | -1661.263 | 1 | 21 | 21 | 43 | 8 | |
| **α3β2β1β2α3** | **0.776929384** | **-98.4** | **-1648.969** | **8** | **23** | **19** | **50** | **9** | 2 |
| α3α3α3β1β2 | 0.74959951 | -111.2 | -1883.27 | 47 | 3 | 2 | 52 | 10 | |
| α3β1β2α3β2 | 0.768982427 | -100.8 | -1704.704 | 18 | 17 | 17 | 52 | 11 | |
| α3α3α3β1β2 | 0.757005931 | -104.8 | -1755.346 | 27 | 14 | 12 | 53 | 12 | |
| **α3β1α3β1β2** | **0.745581393** | **-110.2** | **-1959.395** | **51** | **1** | **3** | **55** | **13** | 10 |
| **α3α3α3β2β1** | **0.756528408** | **-105.1** | **-1683.001** | **29** | **19** | **10** | **58** | **14** | 7 |
| **α3β1α3β2β1** | **0.743622504** | **-107.4** | **-1867.263** | **53** | **6** | **5** | **64** | **15** | 10 |
| α3β1β2α3β2 | 0.781904903 | -94.8 | -1628.559 | 5 | 27 | 32 | 64 | 16 | |
| α3β1α3β2β1 | 0.738593699 | -107.6 | -1877.048 | 60 | 4 | 4 | 68 | 17 | |
| α3α3α3β1β2 | 0.747706278 | -106.2 | -1783.622 | 49 | 13 | 8 | 70 | 18 | |
| α3α3α3β2β1 | 0.752685225 | -99.9 | -1734.166 | 38 | 15 | 18 | 71 | 19 | |
| α3β1β2α3β2 | 0.77890368 | -96 | -1586.335 | 7 | 45 | 24 | 76 | 20 | |
| α3β1β2α3β2 | 0.774357663 | -94.7 | -1602.072 | 11 | 36 | 34 | 81 | 21 | |
| α3β1α3β2β1 | 0.753111003 | -96.1 | -1659.276 | 37 | 22 | 23 | 82 | 22 | |
| α3β1α3β2β1 | 0.742819338 | -101.4 | -1789.493 | 55 | 12 | 16 | 83 | 23 | |
| α3β1α3β2β1 | 0.742463013 | -102.2 | -1716.881 | 57 | 16 | 14 | 87 | 24 | |

| | | | | | | | | | |
|---|---|---|---|---|---|---|---|---|---|
| α3β1β2α3β2 | 0.762403972 | -94.6 | -1618.879 | 21 | 30 | 36.5 | 87 | 25 | |
| α3α3α3β2β1 | 0.751865792 | -95 | -1695.458 | 40 | 18 | 30.5 | 88 | 26 | |
| α3β1β2α3β2 | 0.77471335 | -95.8 | -1554.929 | 10 | 52 | 26 | 88 | 27 | |
| α3α3α3β2β1 | 0.750161078 | -96.9 | -1643.181 | 46 | 24 | 20 | 90 | 28 | |
| α3β1β2α3β2 | 0.757995517 | -94.6 | -1616.385 | 25 | 31 | 36.5 | 92 | 29 | |
| α3α3α3β1β2 | 0.769930939 | -96.2 | -1515.936 | 17 | 60 | 22 | 99 | 30 | |
| α3β1β2α3β2 | 0.760884075 | -95.1 | -1576.559 | 23 | 48 | 29 | 100 | 31 | |
| α3β1β2α3β2 | 0.772805485 | -94.2 | -1556.878 | 14 | 51 | 41.5 | 106 | 32 | |
| **α3β2α3β2β1** | **0.772080348** | **-93.6** | **-1585.683** | **15** | **46** | **45.5** | **106** | **33** | 1 |
| α3α3α3β1β2 | 0.773832944 | -94.5 | -1535.267 | 12 | 57 | 38 | 107 | 34 | |
| **α3α3β1α3β2** | **0.746346204** | **-94.7** | **-1631.43** | **50** | **25** | **34** | **109** | **35** | 3 |
| α3β1α3β1β2 | 0.730239723 | -95.9 | -1662.23 | 65 | 20 | 25 | 110 | 36 | |
| **α3β1β2β2α3** | **0.756582308** | **-94.7** | **-1530.8** | **28** | **58** | **34** | **120** | **37** | 5 |
| α3α3α3β1β2 | 0.768808137 | -94.4 | -1494.306 | 19 | 64 | 39.5 | 122 | 38 | |
| α3β2β1β2α3 | 0.77295397 | -92.7 | -1551.526 | 13 | 53 | 56.5 | 122 | 39 | |
| α3β1α3β1β2 | 0.73708803 | -95.5 | -1601.842 | 63 | 37 | 27 | 127 | 40 | |
| α3β1α3β1β2 | 0.729172241 | -95.3 | -1606.62 | 68 | 33 | 28 | 129 | 41 | |
| α3α3β1α3β2 | 0.771348939 | -92.4 | -1541.793 | 16 | 55 | 59 | 130 | 42 | |
| α3α3β1α3β2 | 0.76605435 | -93.6 | -1493.617 | 20 | 65 | 45.5 | 130 | 43 | |
| α3β1β2β2α3 | 0.758590069 | -93.2 | -1520.286 | 24 | 59 | 49 | 132 | 44 | |
| α3α3α3β1β2 | 0.751587394 | -92.9 | -1595.211 | 41 | 39 | 53.5 | 133 | 45 | |
| α3β1α3β1β2 | 0.727170286 | -95 | -1616.022 | 71 | 32 | 30.5 | 133 | 46 | |
| α3β1α3β1β2 | 0.730211062 | -94.4 | -1619.524 | 66 | 29 | 39.5 | 134 | 47 | |
| α3β1α3β1β2 | 0.729426346 | -94.2 | -1629.897 | 67 | 26 | 41.5 | 134 | 48 | |
| α3β1α3β1β2 | 0.741673411 | -93.5 | -1625.69 | 59 | 28 | 47 | 134 | 49 | |
| α3α3α3β1β2 | 0.755070637 | -93.7 | -1509.603 | 33 | 62 | 44 | 139 | 50 | |
| α3α3α3β1β2 | 0.750162387 | -93.3 | -1560.745 | 45 | 50 | 48 | 143 | 51 | |
| α3α3α3β1β2 | 0.747791151 | -92.7 | -1591.51 | 48 | 41 | 56.5 | 145 | 52 | |
| α3β1α3β1β2 | 0.7373656 | -93 | -1602.607 | 62 | 35 | 51 | 148 | 53 | |
| α3β1α3β2β1 | 0.750995387 | -92.2 | -1587.832 | 44 | 44 | 61.5 | 149 | 54 | |
| α3β1α3β2β1 | 0.742869541 | -92.2 | -1602.632 | 54 | 34 | 61.5 | 149 | 55 | |
| α3α3α3β1β2 | 0.752161168 | -93 | -1508.057 | 39 | 63 | 51 | 153 | 56 | |
| **α3β2β2β1α3** | **0.761033552** | **-91.7** | **-1478.93** | **22** | **66** | **65** | **153** | **57** | 1 |
| α3β1α3β2β1 | 0.737433878 | -92.8 | -1597.43 | 61 | 38 | 55 | 154 | 58 | |
| α3β1α3β1β2 | 0.727455857 | -93.9 | -1589.768 | 70 | 43 | 43 | 156 | 59 | |
| α3β1β2β2α3 | 0.756306667 | -91.6 | -1475.467 | 30 | 67 | 66.5 | 163 | 60 | |
| α3β1β2β2α3 | 0.741848802 | -93 | -1544.561 | 58 | 54 | 51 | 163 | 61 | |
| α3α3α3β1β2 | 0.755225855 | -91.6 | -1464.696 | 32 | 68 | 66.5 | 166 | 62 | |
| **α3β1β2β1α3** | **0.728961325** | **-92.9** | **-1581.732** | **69** | **47** | **53.5** | **169** | **63** | 4 |
| α3α3α3β2β1 | 0.75557732 | -90.9 | -1439.311 | 31 | 71 | 70 | 172 | 64 | |
| α3β1β2β1α3 | 0.711104406 | -92.6 | -1593.73 | 74 | 40 | 58 | 172 | 65 | |
| α3β1β2β2α3 | 0.745398945 | -91.8 | -1539.607 | 52 | 56 | 64 | 172 | 66 | |
| α3β1α3β2β1 | 0.73312626 | -92.3 | -1564.942 | 64 | 49 | 60 | 173 | 67 | |
| α3α3α3β2β1 | 0.754909648 | -91.2 | -1420.613 | 34 | 74 | 68 | 176 | 68 | |
| α3α3α3β1β2 | 0.754213243 | -90.5 | -1429.708 | 36 | 73 | 71 | 180 | 69 | |
| α3β1β2β1α3 | 0.742723547 | -92.1 | -1511.686 | 56 | 61 | 63 | 180 | 70 | |

| | | | | | | | | | |
|---|---|---|---|---|---|---|---|---|---|
| α3β1α3β2β1 | 0.727083067 | -91.1 | -1590.562 | 72 | 42 | 69 | 183 | 71 | |
| α3α3α3β1β2 | 0.7513821 | -89.3 | -1459.74 | 42 | 70 | 72 | 184 | 72 | |
| α3α3α3β2β1 | 0.751044331 | -87.7 | -1390.894 | 43 | 75 | 74 | 192 | 73 | |
| **α3α3β1β1β2** | **0.722332696** | **-88.1** | **-1434.34** | **73** | **72** | **73** | **218** | **74** | 1 |
| α3β1β2β1α3 | 0.702411577 | -85.8 | -1462.193 | 75 | 69 | 75 | 219 | 75 | |

### H: Rankings of Lsa-nAChR2 Dimer Models

Table 6

Ranks of the best unique dimers retrieved from complete pentamers, scored with PRODIGY, ZRANK, and GDOCK. Ranking is based on the sum of the individual scores for PRODIGY, ZRANK, and GDOCK. PRODIGY and ZRANK follow a minimization strategy, meaning a lower score is indicative of a better model. GDOCK follows a maximization strategy, where a higher score is indicative of a better model.

| DIMER | PRODIGY | ZRANK | RANK |
|---|---|---|---|
| α3_β1 | -29.5 | -541.281 | 1 |
| α3_α3 | -23.9 | -437.573 | 2 |
| β2_β1 | -23.8 | -421.116 | 3 |
| β1_α3 | -24 | -402.853 | 4 |
| β2_α3 | -20 | -386.37 | 22 |
| α3_β2 | -21.4 | -383.701 | 28 |
| β1_β2 | -22.9 | -424.312 | 40 |
| β2_β2 | -18.5 | -269.477 | 242 |
| β1_β1 | -14.9 | -259.713 | 373 |

### I: Ranks of all AlphaFold-predicted linked concatemers modelled based on experimental concatemers

Table 7

All generated models of experimentally tested concatenated pentamers. “Concatemer Input” shows the concatemer combination that was modelled in AlphaFold. Concatenated (linked) subunits are indicated using a hyphen (for example α2-α2), while unlinked subunits are indicated using three hyphens (for example α1---β1). The “Concatemer Output” is displayed as a full pentamer in its modelled order, and each linker is given a type of CC (*Counterclockwise*), Cl (*Clockwise*), WdT1 (*Wedge Type 1*), or WdT2 (*Wedge Type 2*). Each “Linker Spacing” is shown as well, indicating if 0, 1, or 2 subunits are wedged between the linked subunits. The AlphaFold metric and Ranking is also shown. The models are grouped by the “Concatemer Input” and new inputs are separated by a blank row.

| Concatemer Input | Concatemer Output | Linker 1 Type | Linker 1 Spacing | Linker 2 Type | Linker 2 Spacing | AlphaFold Score (iptm+ptm) | Rank |
|---|---|---|---|---|---|---|---|
| α2-α2---α1---β1---β2 | α1α2β1β2α2 | WdT1 | 1 | NA | NA | 0.759367646 | 5 |
| α2-α2---α1---β1---β2 | α1α2β1β2α2 | WdT1 | 1 | NA | NA | 0.75741185 | 6 |
| α2-α2---α1---β1---β2 | α1β1β2α2α2 | CC | 0 | NA | NA | 0.749586127 | 8 |
| α2-α2---α1---β1---β2 | α1α2α2β1β2 | CC | 0 | NA | NA | 0.723748202 | 12 |
| α2-α2---α1---β1---β2 | α1α2β1β2α2 | WdT1 | 2 | NA | NA | 0.73235907 | 10 |
| α2-α2---α1---β1---β2 | α1β1β2α2α2 | CC | 0 | NA | NA | 0.716758337 | 14 |
| α2-α2---α1---β1---β2 | α1α2α2β1β2 | CI | 0 | NA | NA | 0.717073664 | 13 |

| | | | | | | | |
|---|---|---|---|---|---|---|---|
| α2-α2---α1---β1---β2 | α1β1α2α2β2 | CC | 0 | NA | NA | 0.751623689 | 7 |
| α2-α2---α1---β1---β2 | α1α2α2β1β2 | CI | 0 | NA | NA | 0.707742755 | 15 |
| α2-α2---α1---β1---β2 | α1α2α2β1β2 | CC | 0 | NA | NA | 0.790243686 | 2 |
| α2-α2---α1---β1---β2 | α1α2β1β2α2 | WdT1 | 1 | NA | NA | 0.787198228 | 3 |
| α2-α2---α1---β1---β2 | α1α2β1β2α2 | WdT1 | 2 | NA | NA | 0.792089972 | 1 |
| α2-α2---α1---β1---β2 | α1α2α2β1β2 | CC | 0 | NA | NA | 0.764282489 | 4 |
| α2-α2---α1---β1---β2 | α1α2β1β2α2 | WdT1 | 1 | NA | NA | 0.732899142 | 9 |
| α2-α2---α1---β1---β2 | α1α2β1β2α2 | WdT1 | 1 | NA | NA | 0.729116521 | 11 |
| α2-α2-β2---α1---β1 | α1β1α2β2α2 | WdT1 | 2 | CC | 0 | 0.719423947 | 12 |
| α2-α2-β2---α1---β1 | α1β1α2α2β2 | CC | 0 | CC | 0 | 0.759785419 | 4 |
| α2-α2-β2---α1---β1 | α1β1α2β2α2 | WdT1 | 2 | CC | 0 | 0.727192025 | 9 |
| α2-α2-β2---α1---β1 | α1α2β2β1α2 | WdT1 | 1 | CC | 0 | 0.699625312 | 13 |
| α2-α2-β2---α1---β1 | α1β1α2α2β2 | CC | 0 | CC | 0 | 0.736974154 | 7 |
| α2-α2-β2---α1---β1 | α1β1α2α2β2 | CC | 0 | CC | 0 | 0.734071238 | 8 |
| α2-α2-β2---α1---β1 | α1β2β1α2α2 | CC | 0 | WdT1 | 1 | 0.674707238 | 14 |
| α2-α2-β2---α1---β1 | α1β1α2α2β2 | CC | 0 | CC | 0 | 0.662826362 | 15 |
| α2-α2-β2---α1---β1 | α1β1β2α2α2 | CC | 0 | WdT2 | 2 | 0.722855721 | 11 |
| α2-α2-β2---α1---β1 | α1β1α2α2β2 | CC | 0 | CC | 0 | 0.776653796 | 1 |
| α2-α2-β2---α1---β1 | α1β2β1α2α2 | CC | 0 | WdT1 | 1 | 0.775338457 | 2 |
| α2-α2-β2---α1---β1 | α1α2β1α2β2 | WdT1 | 2 | WdT1 | 2 | 0.774371198 | 3 |
| α2-α2-β2---α1---β1 | α1α2α2β1β2 | CC | 0 | WdT1 | 1 | 0.752578066 | 5 |
| α2-α2-β2---α1---β1 | α1β2β1α2α2 | CC | 0 | WdT1 | 1 | 0.74716065 | 6 |
| α2-α2-β2---α1---β1 | α1α2β2β1α2 | WdT1 | 2 | CC | 0 | 0.726898232 | 10 |
| α3-α3---β1-β1---β2 | α3β1α3β2β1 | WdT1 | 2 | WdT2 | 1 | 0.757665724 | 11 |
| α3-α3---β1-β1---β2 | α3β1α3β2β1 | WdT1 | 2 | WdT2 | 1 | 0.770522524 | 6 |
| α3-α3---β1-β1---β2 | α3β1α3β2β1 | WdT1 | 1 | WdT1 | 2 | 0.724559496 | 12 |
| α3-α3---β1-β1---β2 | α3α3β1β2β1 | CI | 0 | WdT2 | 1 | 0.757758804 | 9 |
| α3-α3---β1-β1---β2 | α3α3β1β2β1 | CI | 0 | WdT2 | 1 | 0.758445493 | 8 |
| α3-α3---β1-β1---β2 | α3α3β1β2β1 | CI | 0 | WdT2 | 1 | 0.757739949 | 10 |
| α3-α3---β1-β1---β2 | α3α3β1β2β1 | CI | 0 | WdT2 | 1 | 0.720992566 | 13 |
| α3-α3---β1-β1---β2 | α3α3β1β2β1 | CC | 0 | WdT2 | 1 | 0.70943789 | 14 |
| α3-α3---β1-β1---β2 | α3β1α3β1β2 | WdT1 | 2 | WdT1 | 2 | 0.671432237 | 15 |
| α3-α3---β1-β1---β2 | α3α3β1β2β1 | CI | 0 | WdT2 | 1 | 0.802810527 | 2 |
| α3-α3---β1-β1---β2 | α3α3β1β2β1 | CI | 0 | WdT2 | 1 | 0.804260057 | 1 |
| α3-α3---β1-β1---β2 | α3α3β1β2β1 | CI | 0 | WdT2 | 1 | 0.80072816 | 3 |
| α3-α3---β1-β1---β2 | α3α3β1β2β1 | CI | 0 | WdT2 | 1 | 0.775167068 | 5 |
| α3-α3---β1-β1---β2 | α3α3β1β2β1 | CI | 0 | WdT2 | 1 | 0.778778372 | 4 |
| α3-α3---β1-β1---β2 | α3β1α3β1β2 | WdT1 | 1 | WdT1 | 2 | 0.769731165 | 7 |
| α3-α3---β1-β2---β2 | α3β2β2α3β1 | WdT1 | 2 | WdT1 | 2 | 0.801929979 | 2 |
| α3-α3---β1-β2---β2 | α3β1β2α3β2 | WdT1 | 1 | WdT1 | 2 | 0.79969773 | 3 |
| α3-α3---β1-β2---β2 | α3α3β1β2β2 | CI | 0 | WdT2 | 1 | 0.788047266 | 6 |
| α3-α3---β1-β2---β2 | α3β1β2α3β2 | WdT1 | 1 | WdT1 | 2 | 0.770306785 | 10 |
| α3-α3---β1-β2---β2 | α3α3β2β1β2 | CI | 0 | CC | 0 | 0.743936162 | 12 |
| α3-α3---β1-β2---β2 | α3β2β2α3β1 | WdT1 | 2 | WdT1 | 2 | 0.774856563 | 9 |
| α3-α3---β1-β2---β2 | α3α3β1β2β2 | CC | 0 | CC | 0 | 0.539652263 | 15 |
| α3-α3---β1-β2---β2 | α3α3β1β2β2 | CI | 0 | WdT2 | 1 | 0.653875325 | 14 |
| α3-α3---β1-β2---β2 | α3α3β1β2β2 | CC | 0 | WdT2 | 1 | 0.731100438 | 13 |
| α3-α3---β1-β2---β2 | α3β1β2α3β2 | WdT1 | 2 | CC | 0 | 0.802036296 | 1 |
| α3-α3---β1-β2---β2 | α3α3β2β1β2 | CI | 0 | CC | 0 | 0.795191829 | 5 |
| α3-α3---β1-β2---β2 | α3α3β2β1β2 | CI | 0 | CC | 0 | 0.79949012 | 4 |
| α3-α3---β1-β2---β2 | α3β1β2α3β2 | WdT1 | 1 | WdT1 | 2 | 0.775238142 | 8 |

| | | | | | | | |
|---|---|---|---|---|---|---|---|
| α3-α3---β1-β2---β2 | α3β1β2α3β2 | WdT1 | 1 | WdT1 | 2 | 0.769640752 | 11 |
| α3-α3---β1-β2---β2 | α3β2β2α3β1 | WdT1 | 1 | WdT1 | 2 | 0.776645493 | 7 |
| α3-β2---α3---β1---β1 | α3α3β1β2β1 | WdT1 | 1 | NA | NA | 0.764618092 | 6 |
| α3-β2---α3---β1---β1 | α3α3β1β2β1 | WdT1 | 1 | NA | NA | 0.766319373 | 5 |
| α3-β2---α3---β1---β1 | α3α3β1β2β1 | WdT1 | 2 | NA | NA | 0.767358753 | 4 |
| α3-β2---α3---β1---β1 | α3α3β1β2β1 | WdT1 | 1 | NA | NA | 0.729677736 | 13 |
| α3-β2---α3---β1---β1 | α3α3β1β2β1 | WdT1 | 1 | NA | NA | 0.733103386 | 11 |
| α3-β2---α3---β1---β1 | α3α3β1β2β1 | WdT1 | 1 | NA | NA | 0.729737444 | 12 |
| α3-β2---α3---β1---β1 | α3β1α3β2β1 | CC | 0 | NA | NA | 0.671111965 | 15 |
| α3-β2---α3---β1---β1 | α3β1α3β1β2 | CI | 0 | NA | NA | 0.720939939 | 14 |
| α3-β2---α3---β1---β1 | α3β1α3β1β2 | CI | 0 | NA | NA | 0.733684611 | 10 |
| α3-β2---α3---β1---β1 | α3β1α3β1β2 | CI | 0 | NA | NA | 0.778120076 | 3 |
| α3-β2---α3---β1---β1 | α3α3β1β2β1 | WdT1 | 2 | NA | NA | 0.781296363 | 1 |
| α3-β2---α3---β1---β1 | α3β1α3β1β2 | CI | 0 | NA | NA | 0.780039545 | 2 |
| α3-β2---α3---β1---β1 | α3α3β1β2β1 | WdT1 | 2 | NA | NA | 0.735287831 | 9 |
| α3-β2---α3---β1---β1 | α3α3β1β2β1 | WdT1 | 2 | NA | NA | 0.737706222 | 8 |
| α3-β2---α3---β1---β1 | α3β1α3β2β1 | WdT1 | 2 | NA | NA | 0.73971763 | 7 |
| β1-β1---α3-β2---α3 | α3α3β1β1β2 | CC | 0 | WdT1 | 2 | 0.758706075 | 9 |
| β1-β1---α3-β2---α3 | α3β1α3β1β2 | WdT2 | 2 | WdT1 | 1 | 0.789288623 | 4 |
| β1-β1---α3-β2---α3 | α3β1α3β1β2 | WdT1 | 2 | CI | 0 | 0.777233514 | 5 |
| β1-β1---α3-β2---α3 | α3β1α3β1β2 | WdT1 | 2 | CI | 0 | 0.757630708 | 11 |
| β1-β1---α3-β2---α3 | α3β1α3β2β1 | WdT1 | 2 | CC | 0 | 0.745433701 | 13 |
| β1-β1---α3-β2---α3 | α3α3β1β2β1 | WdT2 | 1 | WdT1 | 1 | 0.75527305 | 12 |
| β1-β1---α3-β2---α3 | α3β1α3β1β2 | WdT2 | 2 | CI | 0 | 0.761116227 | 8 |
| β1-β1---α3-β2---α3 | α3β1α3β1β2 | WdT2 | 1 | CI | 0 | 0.642665693 | 15 |
| β1-β1---α3-β2---α3 | α3β1α3β2β1 | WdT2 | 2 | WdT1 | 2 | 0.659530936 | 14 |
| β1-β1---α3-β2---α3 | α3β1α3β2β1 | WdT1 | 2 | WdT1 | 2 | 0.795685361 | 2 |
| β1-β1---α3-β2---α3 | α3β1α3β1β2 | WdT2 | 1 | WdT1 | 1 | 0.792047969 | 3 |
| β1-β1---α3-β2---α3 | α3β1α3β1β2 | WdT1 | 2 | CI | 0 | 0.79678324 | 1 |
| β1-β1---α3-β2---α3 | α3β1α3β1β2 | WdT2 | 1 | WdT1 | 1 | 0.770387404 | 7 |
| β1-β1---α3-β2---α3 | α3β1β1α3β2 | CC | 0 | CC | 0 | 0.758582982 | 10 |
| β1-β1---α3-β2---α3 | α3β1α3β1β2 | WdT2 | 1 | WdT1 | 1 | 0.77101549 | 6 |
| β1-β2-α3---α3---β1 | α3β1α3β1β2 | WdT1 | 2 | Wdt2 | 2 | 0.639404654 | 10 |
| β1-β2-α3---α3---β1 | α3α3β1β2β1 | CI | 0 | WdT1 | 2 | 0.575014209 | 15 |
| β1-β2-α3---α3---β1 | α3α3β1β1β2 | CC | 0 | Wdt1 | 1 | 0.611982663 | 13 |
| β1-β2-α3---α3---β1 | α3β1α3β1β2 | Wdt1 | 2 | CC | 0 | 0.682994145 | 5 |
| β1-β2-α3---α3---β1 | α3α3β1β2β1 | CI | 0 | WdT1 | 2 | 0.625721797 | 11 |
| β1-β2-α3---α3---β1 | α3α3β1β2β1 | CI | 0 | WdT1 | 2 | 0.646915225 | 9 |
| β1-β2-α3---α3---β1 | α3β1β1α3β2 | WdT2 | 1 | CC | 0 | 0.581108542 | 14 |
| β1-β2-α3---α3---β1 | α3β1α3β2β1 | CI | 0 | CI | 0 | 0.654505899 | 8 |
| β1-β2-α3---α3---β1 | α3β1α3β1β2 | WdT1 | 2 | CC | 0 | 0.681354737 | 6 |
| β1-β2-α3---α3---β1 | α3β1α3β2β1 | CI | 0 | CI | 0 | 0.734006636 | 3 |
| β1-β2-α3---α3---β1 | α3α3β1β1β2 | CC | 0 | WdT1 | 1 | 0.723873885 | 4 |
| β1-β2-α3---α3---β1 | α3β1α3β1β2 | CC | 0 | CC | 0 | 0.747129993 | 1 |
| β1-β2-α3---α3---β1 | α3β1α3β1β2 | CC | 0 | CC | 0 | 0.618541643 | 12 |
| β1-β2-α3---α3---β1 | α3α3β1β1β2 | CC | 0 | CC | 0 | 0.743339263 | 2 |
| β1-β2-α3---α3---β1 | α3β1α3β2β1 | CI | 0 | CI | 0 | 0.672191763 | 7 |
| β1-β2---α3---α3---β1 | α3β1α3β2β1 | CI | 0 | NA | NA | 0.769450058 | 4 |
| β1-β2---α3---α3---β1 | α3β1α3β2β1 | WdT2 | 1 | NA | NA | 0.767092679 | 5 |
| β1-β2---α3---α3---β1 | α3β1α3β2β1 | WdT2 | 1 | NA | NA | 0.766406781 | 6 |

| | | | | | | | |
|---|---|---|---|---|---|---|---|
| β1-β2---α3---α3---β1 | α3β1α3β1β2 | WdT1 | 2 | NA | NA | 0.744242975 | 15 |
| β1-β2---α3---α3---β1 | α3β1α3β1β2 | WdT1 | 2 | NA | NA | 0.745007997 | 13 |
| β1-β2---α3---α3---β1 | α3β1α3β1β2 | WdT1 | 2 | NA | NA | 0.744559405 | 14 |
| β1-β2---α3---α3---β1 | α3β1α3β1β2 | WdT1 | 2 | NA | NA | 0.747881607 | 11 |
| β1-β2---α3---α3---β1 | α3β1α3β1β2 | WdT1 | 2 | NA | NA | 0.747577553 | 12 |
| β1-β2---α3---α3---β1 | α3β1α3β1β2 | WdT1 | 2 | NA | NA | 0.75084258 | 10 |
| β1-β2---α3---α3---β1 | α3β1α3β1β2 | WdT1 | 2 | NA | NA | 0.778101516 | 1 |
| β1-β2---α3---α3---β1 | α3β1α3β2β1 | WdT2 | 1 | NA | NA | 0.774738802 | 3 |
| β1-β2---α3---α3---β1 | α3β1α3β2β1 | WdT2 | 1 | NA | NA | 0.77476693 | 2 |
| β1-β2---α3---α3---β1 | α3β1α3β1β2 | WdT1 | 2 | NA | NA | 0.760635413 | 9 |
| β1-β2---α3---α3---β1 | α3β1α3β1β2 | WdT1 | 2 | NA | NA | 0.762377886 | 7 |
| β1-β2---α3---α3---β1 | α3β1α3β1β2 | WdT1 | 2 | NA | NA | 0.761189013 | 8 |
| β2-α1---α2---α2---β1 | α1β1α2β2α2 | WdT1 | 1 | NA | NA | 0.79414609 | 5 |
| β2-α1---α2---α2---β1 | α1α2β2β1α2 | WdT1 | 2 | NA | NA | 0.790107353 | 6 |
| β2-α1---α2---α2---β1 | α1β1α2β2α2 | WdT1 | 1 | NA | NA | 0.794235549 | 4 |
| β2-α1---α2---α2---β1 | α1β1β2α2α2 | WdT1 | 2 | NA | NA | 0.757093489 | 11 |
| β2-α1---α2---α2---β1 | α1α2β1β2α2 | WdT1 | 1 | NA | NA | 0.750864278 | 13 |
| β2-α1---α2---α2---β1 | α1β1β2α2α2 | WdT1 | 2 | NA | NA | 0.757540045 | 10 |
| β2-α1---α2---α2---β1 | α1α2β1β2α2 | WdT1 | 1 | NA | NA | 0.756198078 | 12 |
| β2-α1---α2---α2---β1 | α1β1α2β2α2 | WdT1 | 1 | NA | NA | 0.726346779 | 14 |
| β2-α1---α2---α2---β1 | α1β1β2α2α2 | WdT1 | 2 | NA | NA | 0.715561229 | 15 |
| β2-α1---α2---α2---β1 | α1β1β2α2α2 | WdT1 | 2 | NA | NA | 0.797984694 | 1 |
| β2-α1---α2---α2---β1 | α1β1β2α2α2 | WdT1 | 2 | NA | NA | 0.795562507 | 3 |
| β2-α1---α2---α2---β1 | α1β1β2α2α2 | WdT1 | 2 | NA | NA | 0.795983377 | 2 |
| β2-α1---α2---α2---β1 | α1β1α2β2α2 | WdT1 | 1 | NA | NA | 0.773451523 | 8 |
| β2-α1---α2---α2---β1 | α1α2β1β2α2 | WdT1 | 1 | NA | NA | 0.770686415 | 9 |
| β2-α1---α2---α2---β1 | α1β1β2α2α2 | WdT1 | 2 | NA | NA | 0.773490218 | 7 |
| β2-α3---α3---β1---β1 | α3β1α3β1β2 | WdT1 | 2 | NA | NA | 0.721310673 | 4 |
| β2-α3---α3---β1---β1 | α3β1α3β1β2 | WdT1 | 2 | NA | NA | 0.7120088 | 5 |
| β2-α3---α3---β1---β1 | α3α3β1β2β1 | WdT1 | 1 | NA | NA | 0.674776723 | 7 |
| β2-α3---α3---β1---β1 | α3β1α3β1β2 | WdT1 | 2 | NA | NA | 0.669334499 | 9 |
| β2-α3---α3---β1---β1 | α3α3β2β1β1 | CI | 0 | NA | NA | 0.643140225 | 10 |
| β2-α3---α3---β1---β1 | α3β1α3β1β2 | CC | 0 | NA | NA | 0.685454685 | 6 |
| β2-α3---α3---β1---β1 | α3β1α3β2β1 | CI | 0 | NA | NA | 0.617537728 | 14 |
| β2-α3---α3---β1---β1 | α3β1α3β2β1 | CI | 0 | NA | NA | 0.629482341 | 11 |
| β2-α3---α3---β1---β1 | α3β1α3β2β1 | CI | 0 | NA | NA | 0.628818001 | 12 |
| β2-α3---α3---β1---β1 | α3α3β1β2β1 | WdT1 | 1 | NA | NA | 0.743309356 | 2 |
| β2-α3---α3---β1---β1 | α3α3β1β2β1 | WdT1 | 1 | NA | NA | 0.743472755 | 1 |
| β2-α3---α3---β1---β1 | α3α3β1β2β1 | WdT1 | 1 | NA | NA | 0.739318324 | 3 |
| β2-α3---α3---β1---β1 | α3α3β1β2β1 | WdT1 | 1 | NA | NA | 0.623547657 | 13 |
| β2-α3---α3---β1---β1 | α3α3β1β2β1 | WdT1 | 1 | NA | NA | 0.616433053 | 15 |
| β2-α3---α3---β1---β1 | α3β1α3β1β2 | WdT1 | 2 | NA | NA | 0.671212977 | 8 |
| β2-β1---α3---α3---β1 | α3β1α3β2β1 | WdT1 | 2 | NA | NA | 0.766822097 | 4 |
| β2-β1---α3---α3---β1 | α3α3β1β2β1 | CC | 0 | NA | NA | 0.754565717 | 8 |
| β2-β1---α3---α3---β1 | α3α3β1β2β1 | CC | 0 | NA | NA | 0.754006377 | 9 |
| β2-β1---α3---α3---β1 | α3β1α3β1β2 | WdT1 | 1 | NA | NA | 0.749574806 | 10 |
| β2-β1---α3---α3---β1 | α3α3β1β2β1 | CI | 0 | NA | NA | 0.713341914 | 12 |
| β2-β1---α3---α3---β1 | α3β1α3β1β2 | WdT1 | 1 | NA | NA | 0.740761161 | 11 |
| β2-β1---α3---α3---β1 | α3β1α3β2β1 | WdT1 | 2 | NA | NA | 0.701412153 | 14 |
| β2-β1---α3---α3---β1 | α3β1β1α3β2 | WdT1 | 2 | NA | NA | 0.671800831 | 15 |
| β2-β1---α3---α3---β1 | α3β1α3β2β1 | WdT1 | 2 | NA | NA | 0.702234617 | 13 |

| | | | | | | | |
|---|---|---|---|---|---|---|---|
| β2-β1---α3---α3---β1 | α3β1α3β1β2 | WdT1 | 1 | NA | NA | 0.774681376 | 2 |
| β2-β1---α3---α3---β1 | α3β1α3β2β1 | WdT1 | 2 | NA | NA | 0.778169487 | 1 |
| β2-β1---α3---α3---β1 | α3α3β1β2β1 | CI | 0 | NA | NA | 0.77347791 | 3 |
| β2-β1---α3---α3---β1 | α3β1α3β1β2 | WdT1 | 1 | NA | NA | 0.759638466 | 5 |
| β2-β1---α3---α3---β1 | α3β1α3β1β2 | WdT1 | 1 | NA | NA | 0.754932745 | 7 |
| β2-β1---α3---α3---β1 | α3β1α3β1β2 | WdT1 | 1 | NA | NA | 0.758766681 | 6 |
